# Exopolysaccharide export complex PelBC of *Pseudomonas aeruginosa* attenuates the dynamics of the surrounding outer membrane

**DOI:** 10.64898/2026.08.21.746223

**Authors:** Cristian Rosales-Hernandez, Marius Benedens, Julien Reißmann, Otto Berninghausen, Roland Beckmann, Birgit Strodel, Alexej Kedrov

## Abstract

The opportunistic pathogen *Pseudomonas aeruginosa* ensures its survival by forming mechanically and chemically resistant biofilms, with cationic exopolysaccharide Pel as an abundant constituent of the structural matrix. Despite its biomedical relevance, the mechanisms of Pel synthesis and secretion via the trans-envelope protein machinery are not understood. Here, we examine the structure of the outer membrane export complex PelBC embedded in synthetic nanodiscs and polymer-extracted particles. Both environments preserve the unique architecture of the complex, where the β-barrel PelB is capped with the dodecameric ring of PelC lipoproteins. Cryogenic electron microscopy shows that the polymer-extracted PelB β-barrel is tightly associated with phospholipids and lipid A molecules, and the membrane-facing PelC ring may stabilize lipids of the periplasmic leaflet in defined positions. All-atom molecular dynamics simulation of PelBC in the asymmetric outer membrane of *P. aeruginosa* corroborate the structural findings and visualize how the essential C-terminal helix of PelC forms multiple electrostatic contacts with the periplasmic leaflet of the outer membrane. Those interactions reduce the lateral mobility of the lipids, stabilize the position of the ring at the interface and may guide folding and assembly of the polysaccharide export machinery.

**Highlights:**

- The PelBC complex is visualized in nanodiscs and polymer-extracted particles
- The architecture of PelBC is not affected by the chosen membrane mimetics
- Structure-based molecular dynamics simulations validate PelBC:lipid interactions
- Lipid mobility in the outer membrane is hindered by the embedded PelBC complex

## Introduction

Biofilms are microbial communities that may consist of one or multiple species embedded within a self-produced extracellular matrix, often formed at solid surfaces or liquid-air interfaces [1]. Biofilms are linked to a broad range of chronic and recurrent infections, as they contribute to the survival of microbial pathogens in diverse environments [2]. The mechanical stability and resilience of biofilms are mainly attributed to the extracellular polymeric substance (EPS) matrix, which is composed of proteins, extracellular DNA, lipids or surfactants, water, and exopolysaccharides [3]. *Pseudomonas aeruginosa*, an opportunistic human pathogen and a global health threat, produces several exopolysaccharides, including alginate, Psl and Pel [4]. Biogenesis of Pel, a cationic N-acetylgalactosamine-based polymer found in pellicles,i.e. biofilms at the air-water interface, requires a set of proteins encoded within the *pelABCDEFG* operon [5]. The cell envelope-spanning machinery mediates synthesis and transport of the polysaccharide across both membranes of *P. aeruginosa* (Figure 1A) [4]. The coupled synthesis and transport across the inner membrane are facilitated by the membrane-embedded complex PelDEG and the cytosolic gylcosyltransferase PelF [6]. Once exposed to the periplasm, the polysaccharide is partially deacetylated by the enzyme PelA, and the final export step is mediated by the PelBC complex at the outer membrane [7].

**Figure 1.**
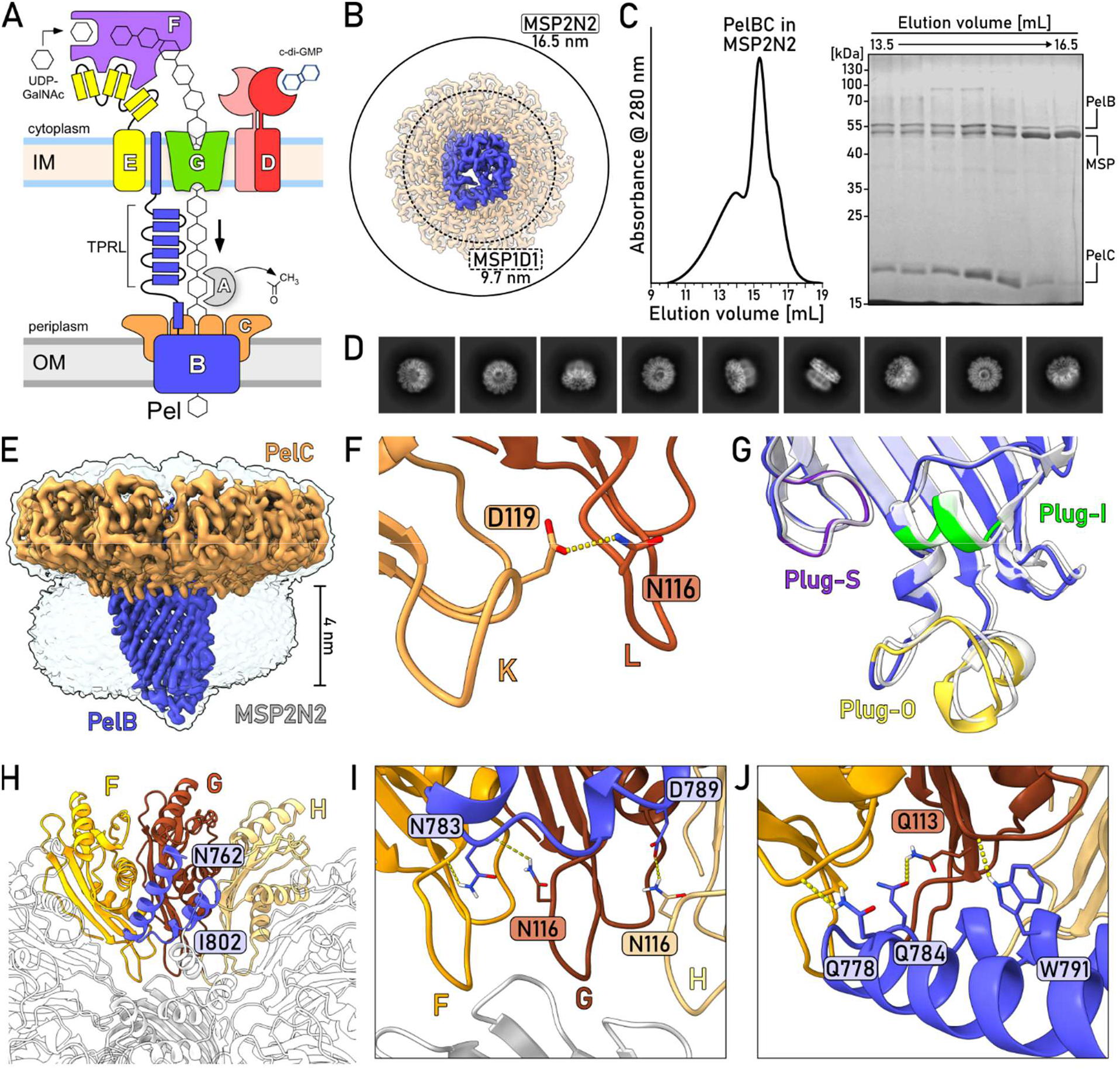
Architecture of PelBC in MSP2N2-based nanodiscs. **(A)** Schematics of the Pel exopolysaccharide secretion system spanning the inner (IM) and the outer membrane (OM) of *P. aeruginosa*. The individual subunits are indicated with their corresponding letters, e.g. “B” for PelB. **(B)** PelBC dimensions in comparison to the expected outer diameters of nanodiscs formed by MSP1D1 (inner ring, dashed line) and MSP2N2 scaffold proteins (outer ring, solid line). **(C)** Size exclusion chromatography profile and the corresponding SDS-PAGE of PelBC reconstituted in MSP2N2-based nanodisc. The protein bands are indicated. **(D)** Representative 2D classes from single-particle cryo-EM analysis of PelBC reconstituted into MSP2N2-based nanodiscs. **(E)** Final 3D reconstruction of the PelBC complex in MSP2N2-based nanodisc. Two different signal levels are shown to visualize PelBC only (low levels; colored) and the surrounding membrane density (high levels; white). **(F)** The interaction between Asp-119 and Asn-116 of proximate PelC subunits (on example of PelC subunits K and L). **(G)** Configurations of the extracellular loops Plug-I (green), Plug-O (yellow), and Plug-S (violet) of PelB resolved in MSP2N2-based nanodiscs. The conformation of PelBC in MSP1D1-nanodisc is shown in white (PDB ID 9H80). **(H)** Residues 762-802 form a helical bundle at the N-terminal end of PelB (blue) that is docked against PelC subunits F, G and H. **(I and J)** Interactions of the N-terminal domain of PelB (blue) with PelC subunits F, G, and H.

The C-terminal domain of PelB forms a 16-stranded β-barrel that serves for exporting the polysaccharide across the outer membrane, and its architecture is homologous to BcsD and PgaA, the export proteins for bacterial cellulose and poly-N-acetylglucosamine (PNAG) [8, 9]. Apart from the similar structure of the β-barrel, all those proteins contain N-terminal helical scaffolds exposed to the periplasm. The scaffold of PelB is sufficiently long to span the periplasm, and it is likely anchored in the inner membrane by a single transmembrane helix [10]. Thus, it may bridge the inner membrane biosynthetic machinery PelDEFG with the outer membrane export pore and build a continuous passage for the polysaccharide. Another distinctive feature of the Pel export machinery is the outer membrane lipoprotein PelC that assembles into a ring-shaped complex composed of twelve subunits [7]. Though its role remains elusive, deletion of PelC abolishes the exopolysaccharide export, and several mutations impair biofilm formation [7, 11].

Recently, we employed cryogenic electron microscopy (cryo-EM) to reveal the structure of PelBC in the lipid environment, and combined molecular dynamics (MD) simulations and single-channel conductivity analysis to probe the conformational dynamics of PelB [10]. The structure confirmed that the complex consists of a β-barrel PelB capped on the periplasmic side with a dodecameric ring of PelC. A repertoire of interactions at the barrel:ring interface stabilized the assembly. The periplasm-facing loops at the rim of the PelB barrel built electrostatic interactions with Asp-119 of multiple PelC subunits, and additional contacts were formed via the N-terminal helical region of PelB. The acyl chains conjugated at the terminal cysteine residue of PelC (position 19 prior to the signal peptide cleavage) and the residues Trp-149 served to anchor the ring into the lipid leaflet. Lipid-like densities were observed around the β-barrel likely stabilized via electrostatic and hydrophobic contacts with the protein surface, and the complementary MD simulation of the membrane-embedded PelB suggested that the lipid A molecules may tightly bind to the cationic clusters at the extracellular side of PelB.

The architecture of PelBC was resolved in the relatively small MSP1D1 lipid-based nanodisc, limiting the membrane diameter to 8 nm [12]. While the provided area was sufficient to accommodate the PelB β-barrel of 4 nm and the surrounding anchors of PelC, the diameter of the PelC ring (above 12 nm) exceeded the nanodisc dimensions (Figure 1B), so the reconstituted system did not allow studying the protein:lipid interactions at the interface. Furthermore, detergent extraction of PelBC prior the nanodisc reconstitution might perturb native lipid interactions or affect the complex architecture, and the origin of the lipid-like densities observed near the β-barrel cannot yet be assigned unambiguously, as these lipids may derive from the native membrane or may have been introduced during the reconstitution procedure. To address those issues, we examine here the architecture of the PelBC complex in the significantly larger MSP2N2 nanodisc, and in polymer-based lipid particles, which allowed us to bypass the detergent solubilization step. We confirm that the overall PelBC architecture is preserved across distinct membrane-mimetic environments, but also resolve additional features of the PelB N-terminal region, and a diversity of lipids, including lipid A, associated with the protein complex. All-atomistic molecular dynamic simulations of PelBC in native-like asymmetric membranes confirm the extensive network of protein:lipid interactions and suggest that the PelC ring builds a broad contact area with the periplasmic leaflet, which is potentially required for PelBC assembly.

## Results

### The architecture of the reconstituted PelBC is not affected by the nanodisc dimensions

To assess whether the previously resolved structure was not affected by the dimensions of the MSP1D1-based nanodisc, we set out to reconstitute and visualize PelBC in a substantially larger scaffold MSP2N2, which typically forms nanodiscs of ~17 nm diameter (Figure 1B) [13]. Size exclusion chromatography (SEC) of the reconstitution reaction manifested a major peak at 15.5 mL that correlated with the expected mass of a PelBC complex within a nanodisc (350-400 kDa; Figure 1C). The leading SEC shoulder at ~14 mL potentially contained nanodiscs with two copies of the PelBC complex in opposing topologies, and the tail shoulder at ~16 mL contained predominantly the scaffold protein MSP2N2 (Figure 1C), likely as a part of lipid-only nanodiscs.

The cryo-EM and single-particle analysis of the major elution peak showed a broad distribution of angular views of PelBC (Figure 1D), and the resulting 3D reconstruction revealed the architecture of the complex at global resolution of 2.6 Å (Figure 1E and Suppl. Figures 1, 2A and B). Due to the large area of the lipid bilayer, the MSP2N2-based reconstruction provided a clear view of the orientation of PelBC relative to the membrane. The PelC ring lies nearly parallel to the membrane plane, whereas the PelB β-barrel is positioned slightly tilted within the bilayer. The resolved PelBC architecture appeared nearly identical to the previous model (RMSD of 0.339 Å, all-atom; Suppl. Figure 3A) [10], so the dimensions of the smaller nanodisc did not substantially influence the complex. Furthermore, several refinements of the structural model were possible due to improved local resolution. In particular, the position of the flexible D-loop of PelC subunit K (PelC_K_, residues 116-121) was unambiguously modelled, as the MSP2N2-based map manifested a continuous density in this region (Suppl. Figure 3B). In the earlier model, Asp-119 of PelC_K_ was proposed to interact with a periplasmic loop of PelB and serve to anchor the ring to the β-barrel [10]. The new reconstruction shows a minor displacement within PelC_K_, so Asp-119 appears within approx. 2.5 Å from Asn-116 of the neighbouring subunit PelC_L_, suggesting that the hydrogen bond between the residues stabilizes the internal architecture of the PelC ring (Figure 1F and Suppl. Figure 3C). The pore-facing loops of PelC subunits (residues 115-121) were further stabilized by intermolecular hydrogen bonds formed by Gln-46 and Gln-49, thus providing a rigid interface for docking the periplasmic domain of PelB (Suppl. Figure 3D, also described below). The PelB β-barrel was uniformly present in a closed conformation, sealed by the extracellular loops Plug-I (residues 997-1002), Plug-O (residues 1077-1090) and Plug-S (residues 1171-1177) (Figure 1G, Suppl. Figures 4A and B).

Built of multiple TPR-like (TPRL) helical domains, the N-terminal domain of PelB may serve as a physical bridge for the polysaccharide chain crossing the periplasm. The PelB construct designed for structural analysis contained three TPRL domains prior to the membrane-embedded β-barrel. Whereas the local resolution was previously insufficient to model the constituting residues 762 to 802, the segment has been fully resolved in the new reconstruction, possibly due to differences in particle orientation between the cryo-EM datasets (Figure 1H, Suppl. Figure 2A and 5A). The TPRL domains formed contacts via Asn-783 and Asp-789 to Asn-116 of the proximate PelC subunits F, G, and H (Figure 1I). In addition, Gln-778, Gln-784, and Trp-791 were positioned close to Gln-113 and backbone atoms of the central PelC loop (subunit F and G; Figure 1J). The N-terminal domain of PelB was aligned with a cluster of anionic residues within the β-barrel and the putative gating loop Plug-S, thus suggesting a route taken by the exopolysaccharide across the export complex (Suppl. Figure 5B and 5C).

### Polymer-extracted PelBC retains its core architecture

The comparative analysis of PelBC in MSP1D1- and MSP2N2-based nanodiscs has provided a consolidated view of the protein complex in the lipid environment and indicated that its overall organization is not substantially influenced by the dimensions of the nanodiscs. Next to the detailed reconstruction of the PelBC complex, the analysis revealed a repertoire of rod-shaped densities, which were assigned to acyl chains, and their positions correlated between the studied samples [10] (Suppl. Figure 6A). Many of these densities corresponded to the lipid anchors attached to the N-terminal cysteine of each PelC subunit (Suppl. Figure 6B), while others reflected lipid molecules stabilized in the vicinity of the PelB β-barrel. Those lipids were potentially introduced along the reconstitution procedure, but they could also originate from the bacterial membrane, being co-purified with the protein complex. Indeed, several lipid molecules were docked in narrow pockets between the β-barrel and the anchors of the PelC ring, so they may have adopted their positions during the PelB-PelC assembly in the cell. Furthermore, the densities visualized within the extracellular leaflet appeared rather bulky for phospholipids [10], suggesting that they may correspond to lipid A, an essential component of bacterial outer membranes. Lipid A molecules can stably bind proteins via hydrophobic acyl chains, but also electrostatic interactions between the phosphate moieties and cationic amino acids, so they have been resolved in multiple crystal and cryo-EM structures of bacterial outer membrane proteins (OMPs) [14–16].

To assess PelBC:lipid interactions formed in the bacterial membrane, we set out to isolate the protein complex with its proximate environment, so instead of using detergent micelles for the extraction, we turned our attention to amphipathic co-polymers [17]. Co-polymers, such as styrene maleic acid (SMA) and its derivatives, have been extensively used for isolation of bacterial and eukaryotic membrane proteins, and a few polymer-extracted OMPs have been studied by cryo-EM, exemplified by OmpF, PagP and the assembled BAM machinery [18–20]. However, SMA-type polymers are often sensitive to changes in pH, ionic strength, and presence of divalent cations. Thus, we have focused on the recently developed cycloalkane-modified amphipathic polymers (marketed as Ultrasolute amphipol 18, further referred to as UApol18, Suppl. Figure 7A) which manifest advanced stability over a broad range of conditions [21, 22]. Though the feasibility of UApol18 application to OMPs and their complexes was not previously assessed, the polymer has been used to determine structures of inner membrane proteins, such as the efflux pump AcrB [23] and the ABC transporter MsbA [24]. Importantly, with the estimated hydrodynamic diameter of 16 nm [25], UApol18-based nanodiscs should be sufficiently large to embed the β-barrel of PelB and to capture the membrane under the PelC ring.

Both PelB and PelC were efficiently extracted from the cellular membranes upon incubation with UApol18, and immobilized metal affinity chromatography (IMAC) targeted for the polyhistidine-tagged PelB subunit led to isolation of both proteins (Figure 2A). Mass photometry recordings showed a major peak at ~350 kDa that matched closely the mass of PelBC in detergent micelles (Figure 2B), and negative-stain electron microscopy visualized abundant doughnut-shaped particles expected for the PelC ring (Suppl. Figure 2B), so we concluded that the complex retained its integrity. Hence, despite several distinct impurities seen in SDS-PAGE (Figure 2A), the sample was directly subjected to cryo-EM analysis, which confirmed presence of the assembled PelBC (Figure 2C and Suppl. Figure 8). The resulting 3D reconstruction and the structural model of the polymer-extracted complex was nearly identical to the structures obtained in the lipid-based MSP nanodiscs (RMSD to 9H80: 0.342 Å, all-atoms; Figures 2D and E), suggesting that the architecture was not substantially affected by the detergent-based isolation and following reconstitution procedure. Notably, the polymer-extracted complex manifested once again a uniformly closed conformation of the β-barrel, which likely represent the idle state of the export machinery in absence of the exopolysaccharide (Suppl. Figure 4C and D). Interestingly though, the resolved periplasmic domain of PelB was limited to the stalk helix (residues 862-879), which could be possibly attributed to preferential orientation of the particles seen in 2D classification (Suppl. Figure 9A).

**Figure 2.**
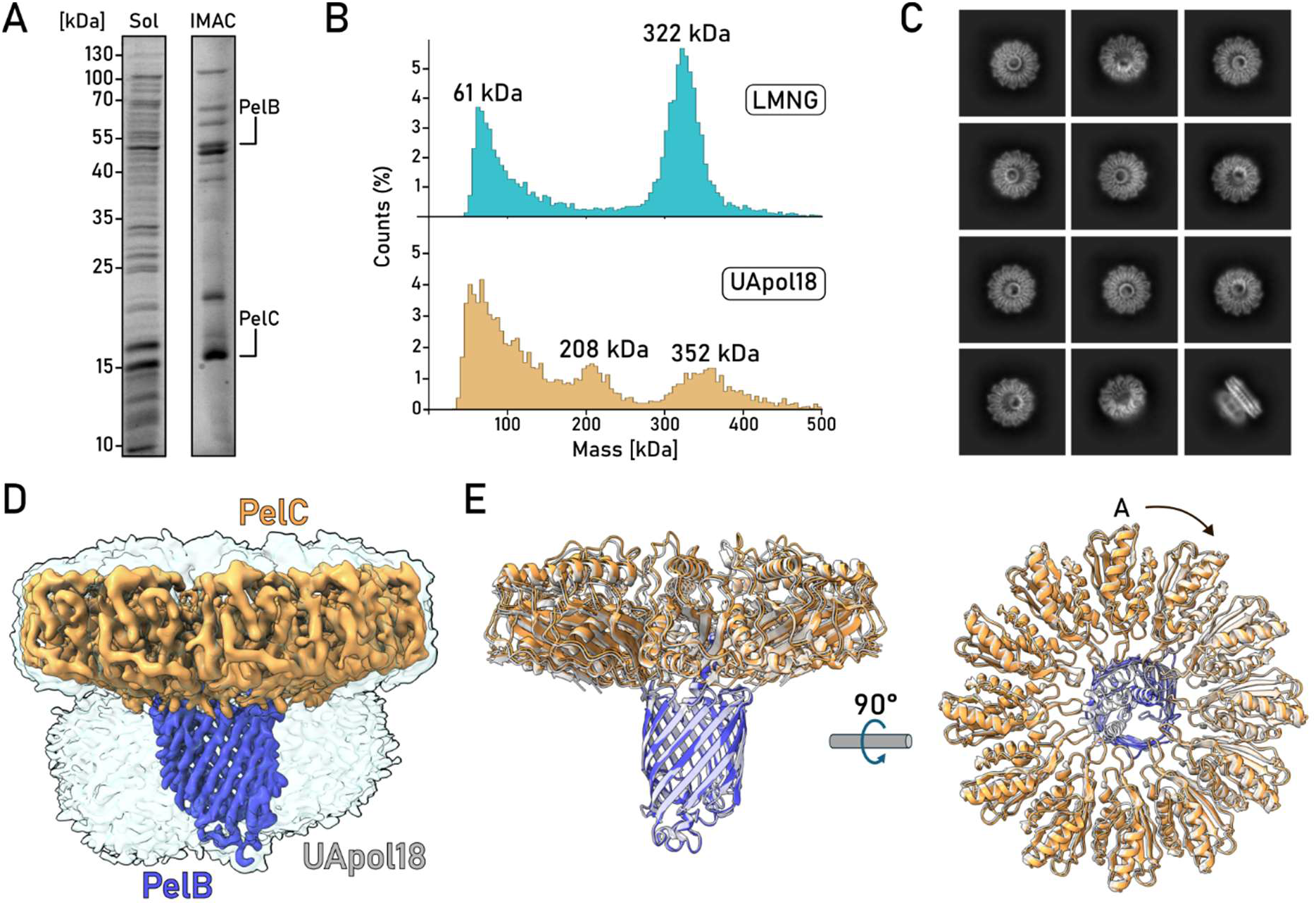
Architecture of PelBC in UApol18-based nanodiscs. **(A)** SDS-PAGE of the UApol18-solubilized material (total membrane extract) and the IMAC elution fraction. The bands corresponding to PelB and PelC are indicated. **(B)** Distribution of particle masses measured for the purified PelBC complex in LMNG (top) and UApol18 (bottom) measured by single-molecule mass photometry. **(C)** Representative 2D classes from single-particle cryo-EM analysis of the PelBC complex in UApol18. **(D)** Final 3D reconstruction of the PelBC complex in UApol18. Two different signal levels are shown to visualize PelBC only (low levels; colored) and the surrounding membrane density (high levels; white) **(E)** Overlay of the structural models of PelBC resolved in UApol18 (colored) and MSP1D1-based nanodiscs (white; PDB ID 9H80).

### Cryo-EM analysis reveals extensive PelBC:lipid interactions

The measured mass of PelBC in UApol18 (~350 kDa; Figure 2B) substantially exceeded the calculated molecular weight of the protein alone (250 kDa), and cryo-EM reconstruction revealed a broad micelle-like coat around the β-barrel. Thus, the complex was likely extracted with a fraction of the bacterial outer membrane (Figure 2D). The acyl chain anchors of PelC subunits could be readily seen within the “coat” (Suppl. Figure 6A and B), but also several free-standing densities near the β-barrel were resolved, implying that the outer membrane lipids were docked and co-purified at these sites (further referred as annular lipids [26]) (Figure 3A). Two lipid molecules were resolved near the lipid anchor of PelC_E_ and PelC_D_ (Figure 3B), and another lipid molecule was located in proximity to PelC_I_, with a density sufficient to trace both acyl chains for the first four to five carbons (Figure 3C). While bulky aromatic and apolar residues of PelB β-barrel formed clefts for the acyl chains, electrostatic interactions at the membrane interface were mediated by PelC. Here, Arg-151 of PelC_C_ and PelC_I_, a residue implicated in biofilm formation [7], were reaching the phosphates of the proximate annular lipids. Proximate Asp-119 were oriented to the head groups, suggesting that phosphatidylethanolamine (PE) lipids would be preferred at those positions due to electrostatic interactions (Suppl. Figure 10). Multiple lipid densities were also found beyond the PelB:PelC interface, towards the periphery of the PelC ring (further referred as *non-annular lipids*), and their positions correlated well between MSP-and UApol18-based samples (Figure 3A). Those lipids must be transiently stabilized via electrostatic interactions with PelC, with an example of PelC_D_ Arg-146 oriented towards the membrane, so it could form a bond with a phosphate moiety of a lipid (Figure 3B). While PelC exposed multiple charged residues towards the periplasmic leaflet, where a pattern of densities was observed in cryo-EM maps (Figure 3A), further analysis of protein:lipid interactions was hindered by the lower resolution within the dynamic lipid interface.

**Figure 3.**
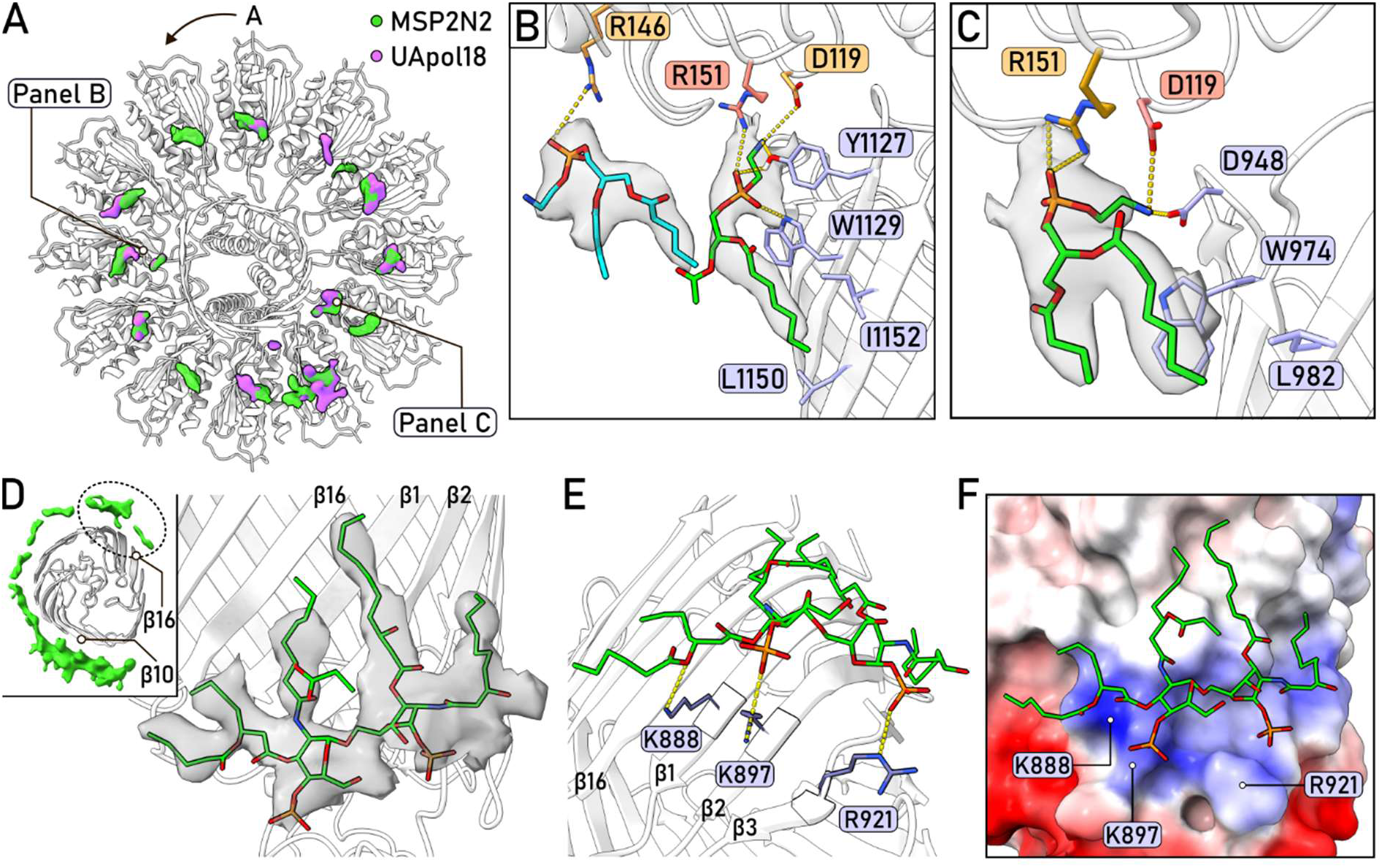
Protein:lipid interactions resolved by cryo-EM for the UApol18-extracted PelBC. **(A)** Comparison of the lipid densities under the PelC ring resolved in MSP2N2- and UApol18-based reconstructions (shown in green and purple, respectively). **(B)** Lipid interactions with residues of PelB (blue) and PelC (residues of subunit C shown in salmon; D in orange). Non-annular PE lipid is shown in green, annular PE is shown in cyan. **(C)** Interactions of a non-annular PE lipid with PelB (blue) and PelC subunits H (residues shown in orange) and I (salmon). **(D)** Electron density at the extracellular side of the PelB β-barrel with a docked fragment of lipid A molecule. **(E)** Interactions of the positively charged PelB residues with the lipid A molecule. **(F)** Electrostatic potential of PelB visualizes a positively charged interface for binding lipid A.

While MSP-based nanodiscs cannot recapitulate the natural asymmetry of the bacterial outer membrane, UApol18 potentially extracted PelBC-bound lipids from both leaflets. Indeed, the polymer-based reconstruction revealed a prominent density within the extracellular leaflet that formed close contact with PelB β-strands β1, β2, β3 and β16 (Figure 3D) and manifested five “tails” within the membrane. The shape and the dimensions matched well those of a lipid A molecule, the main component of the outer membrane that contains up to six acyl chains. The modelled lipid A molecule was positioned in close proximity to Lys-888, Lys-897 and Arg-921 of PelB, so its phosphate groups appeared within 3-5 Å distance to the positively charged residues, and additional contacts were formed via the oxygen atoms within the lipid A (Figure 3E and F). We also observed a broad density on the opposite side of the β-barrel, near the strands β10 to β12 (Figure 3D, inset), which could originate from one or more lipid A molecules, which however remained too dynamic to be resolved in detail. Binding of lipid A molecules at those sites of PelB via electrostatic interactions was observed in previous simulations [10], and the cryo-EM reconstruction in the polymer-based nanodiscs further corroborates that interaction model.

### Molecular dynamics simulations of PelBC in the outer membrane

Cryo-EM analysis of PelBC in the membrane environments has revealed a diversity of protein:lipid interactions formed across the asymmetric bilayer. The visualized anchors of the PelC lipoproteins are essential for the complex assembly [7], while the phospholipids and lipid A molecules surrounding the β-barrel of PelB may contribute to its stability. Notably, the contacts of the PelC ring with the periplasm-facing membrane leaflet seen in both MSP2N2- and UApol18-based samples suggested that there might be preferential, though likely dynamic interactions between the protein and lipid headgroups at the periphery of the complex, which are challenging to address via structural analysis. Indeed, a pattern of densities at the membrane interface was observed (Figure 3A), matching positions of PelC subunits, which could reflect lipid head groups in contact with the protein; however, the interpretation based solely on cryo-EM data remained ambiguous.

To obtain a complementary view, we performed all-atom molecular dynamics (MD) simulations of the PelBC:membrane system. Our previous study has already addressed the conformational dynamics of PelB in symmetric and asymmetric membranes and revealed binding sites for lipid A molecules, in agreement with the structural data [10]. With the present simulations, we aimed to expand the analysis to the whole PelBC complex embedded into a physiologically relevant asymmetric membrane, focusing the attention on the protein:lipid interaction network. As the PelBC reconstructions appeared nearly identical across the tested mimetics, the MSP1D1-based model was chosen for the simulations. The protein complex was embedded into an asymmetric model of the *P. aeruginosa* outer membrane with lateral dimensions of 18.59 × 18.59 nm^2^ (Figure 4A). The periplasmic leaflet was composed of DPPE, DPPG, and DOPE, while the outer leaflet consisted of lipid A [27]. To model the post-translational triacylation of the lipoprotein PelC, lipid anchors were covalently bound to Cys-19 of each PelC subunit. The ion concentration was set to 25 mM NaCl to approximate the growth conditions of *P. aeruginosa* at the air-water interface. The initial position of the complex was selected so that the β-barrel of PelB and the acyl chain anchors of PelC were inserted into the membrane, while the PelC ring did not form contacts with the periplasmic lipid leaflet (Suppl. Figure 11). However, along each simulation, the complex spontaneously shifted towards the membrane by approx. 0.85 nm, so PelC subunits closely approached lipid head groups. The ring positioned itself parallel to the membrane plane, thus matching the cryo-EM reconstructions. Accordingly, the β-barrel showed a tilt of approx. 13°, while it was oriented nearly orthogonal to the membrane in the previous modelling (Suppl. Figure 12) [10].

**Figure 4.**
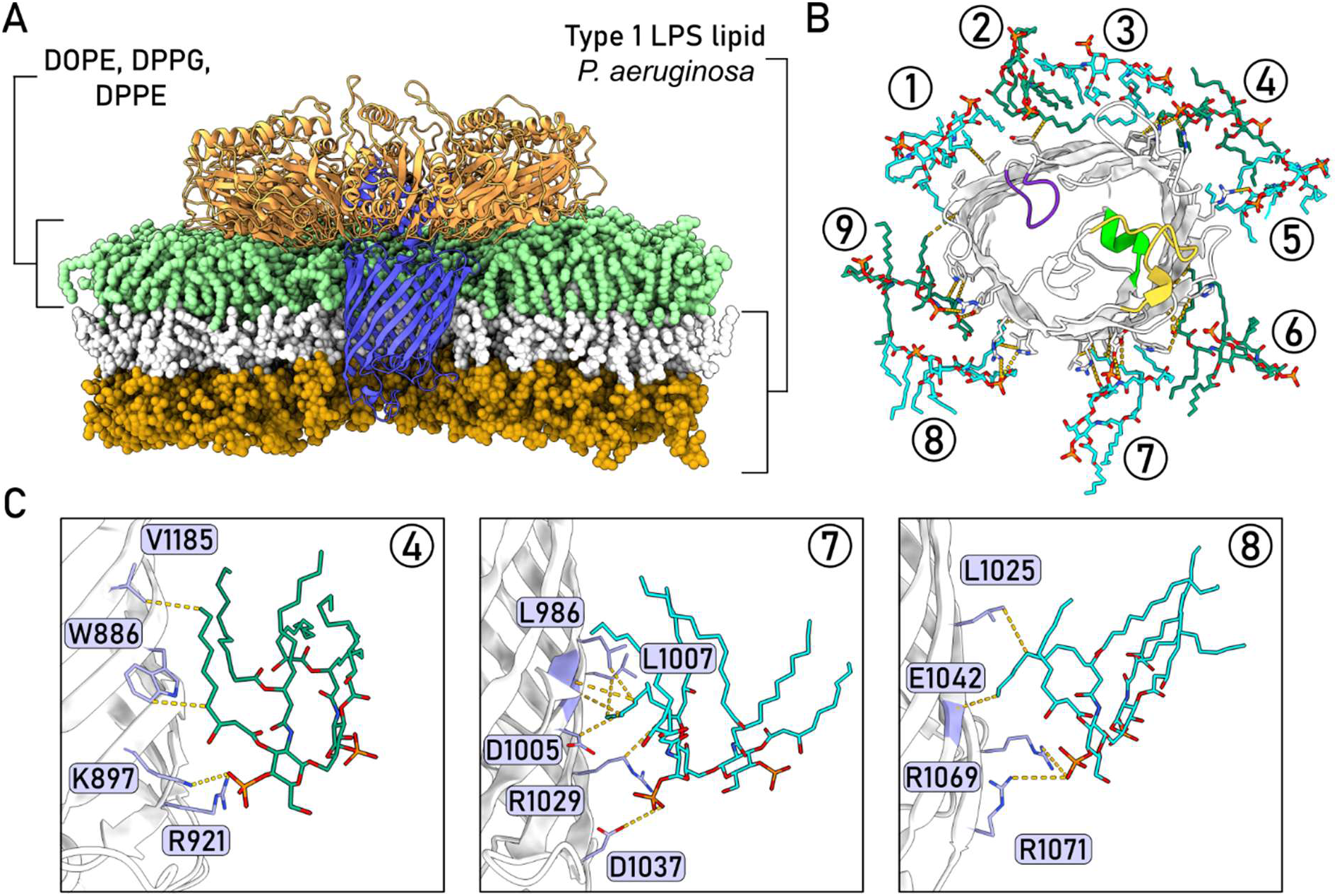
PelB builds multiple interactions with lipid A molecules of the outer membrane. **(A)** Visualization of the PelBC complex in the asymmetric outer membrane of *P. aeruginosa* used in the MD simulation. The lipids of the periplasmic leaflet are shown in light green, the lipid A moieties of lipopolysaccharide (LPS) in white, and the core oligosaccharide in sand yellow. **(B)** A representative distribution of lipid A molecules at the extracellular side of the PelB β-barrel, as observed in the MD simulation. The numbered Lipid A molecules are shown in alternating green and cyan. **(C)** Examples of interactions formed between PelB and lipid A molecules. The lipids are numbered (in circles) accordingly to panel B.

The PelBC complex retained its overall structural integrity across all simulations, with conformational dynamics confined to local fluctuations around the native fold (Suppl. Figure 13). As previously observed for PelB alone, the extracellular loops Plug-O and Plug-S were relatively mobile, while the structurally essential α-helix Plug-I inside the β-barrel showed no displacement. Notably, we observed now enhanced dynamics of the loop between β-strands 3 and 4 (residues 924-936), which remained static in the earlier analysis. The loop flexibility could be influenced by the specific ionic strength (25 mM vs. 150 mM NaCl in earlier simulations) or the geometry of the β-barrel differing between the simulations, and so altered contacts with the membrane. Although the role of the Plug-S as a gate for Pel export is supported by several lines of evidence, such as its alignment with the TPRL domains and experimentally observed fluctuations between closed and open states [10], the exact route of the polysaccharide chain remains to be elucidated. Consequently, the observed conformational dynamics at the PelBC exit warrant consideration in future dedicated studies.

UApol18-based reconstruction suggested that lipid A molecules tightly associate with the PelB β-barrel, so we used the MD simulation to trace those interactions. Being negatively charged inside the barrel and at the extracellular loops, PelB exposes multiple lysines and arginines at the extracellular rim, which may serve as lipid binding sites. Indeed, nine to ten lipid A molecules were found surrounding the barrel along the simulations (Figure 4B), and the measured frequency of contacts with specific PelB residues allowed to map the interactions (Suppl. Figure 14). In agreement with the structural data, Lys-897 and Arg-921 formed stable contacts with one lipid A molecule that remained bound to the β-barrel over the course of the simulation (Figure 4C, lipid #4).

These interactions with the phosphate group were further supported by docking against Trp-886 and Val-1185 in the depth of the membrane. Two copies of lipid A were bound on the opposing side of the β-barrel rim, with their phosphate groups oriented towards Arg-1029, Arg-1069, and Arg-1071 (Figure 4C, lipids #7 and #8), thus corroborating our hypothesis regarding the origin of the broad density observed in the cryo-EM map (Figure 3D, inset).

### The ring of PelC subunits modulates the dynamics of the proximate lipids

MD simulations of PelBC validated that the complex was stably anchored within the native-like membrane, and the PelC ring closely approached the periplasmic leaflet. To characterize the lipid dynamics and protein:lipid interactions there, we analyzed the lipid dwell times, as these should reveal positions of PelBC-bound and thus less mobile lipids (Figure 5B and Suppl. Figure 15). Indeed, the two-dimensional maps suggest non-uniform lipid dynamics around PelBC, as it could be divided into three distinct areas: (i) lipid-free space occupied either by the β-barrel or the “belt” of the PelC anchors (white areas of the map), (ii) annular lipids in contact with PelB, shielded by the PelC anchors and the intercalating Trp-149 residues (marked with a bold solid circle), and (iii) non-annular lipids beyond the anchors, but underneath the PelC ring. In accordance with the cryo-EM reconstructions, the annular lipid molecules docked at the surface of PelB and shielded by PelC anchors (subunits H, I and J) remained stably bound in their positions. Interestingly, a long-term interaction at subunit PelC_I_ was repeatedly observed for a DPPE molecule, as its amine group formed hydrogen bonds with the proximate aspartate residues of PelC and PelB (Figures 3C and 5A; Suppl. Figure 16A). When a DPPG molecule was docked at the same position, it was expelled and substituted by PE within 300 ns of the simulation (Suppl. Figure 16B), thus highlighting the role of electrostatic interactions at the interface.

Arranged into a ring with a slightly conical shape, the PelC subunits came into the closest contact with the annular lipids near the rim of the PelB β-barrel. However, the surrounding non-annular lipids were also affected by the membrane-exposed N-terminal β-strand/loop (residues 20-27) and the C-terminal helix (residues 156-172) of each PelC subunit. The lipid dwell time map shows a broad low-mobility area at the periphery of PelC, where a pattern of densities was observed in the cryo-EM reconstructions (Figure 3A). To assess the protein:lipid interactions there, we analyzed the frequencies of lipid contacts formed by residues of the individual PelC subunits. Because the ring was aligned with the membrane plane, the pattern of interactions was similar among the subunits (Figure 5C and D; Suppl. Figures 17 and 18). As expected, the charged residues within the membrane-exposed domains, such as Arg-146, Lys-160, Arg-163, Glu-164, Asp-168 and Arg-170, formed the most extensive contacts with the lipid headgroups (Figure 5A and Suppl. Figure 19). This remarkable agreement between the structural data and the simulations indicates that lipid diffusion within the membrane is attenuated via electrostatic interactions with PelC lipoproteins.

**Figure 5.**
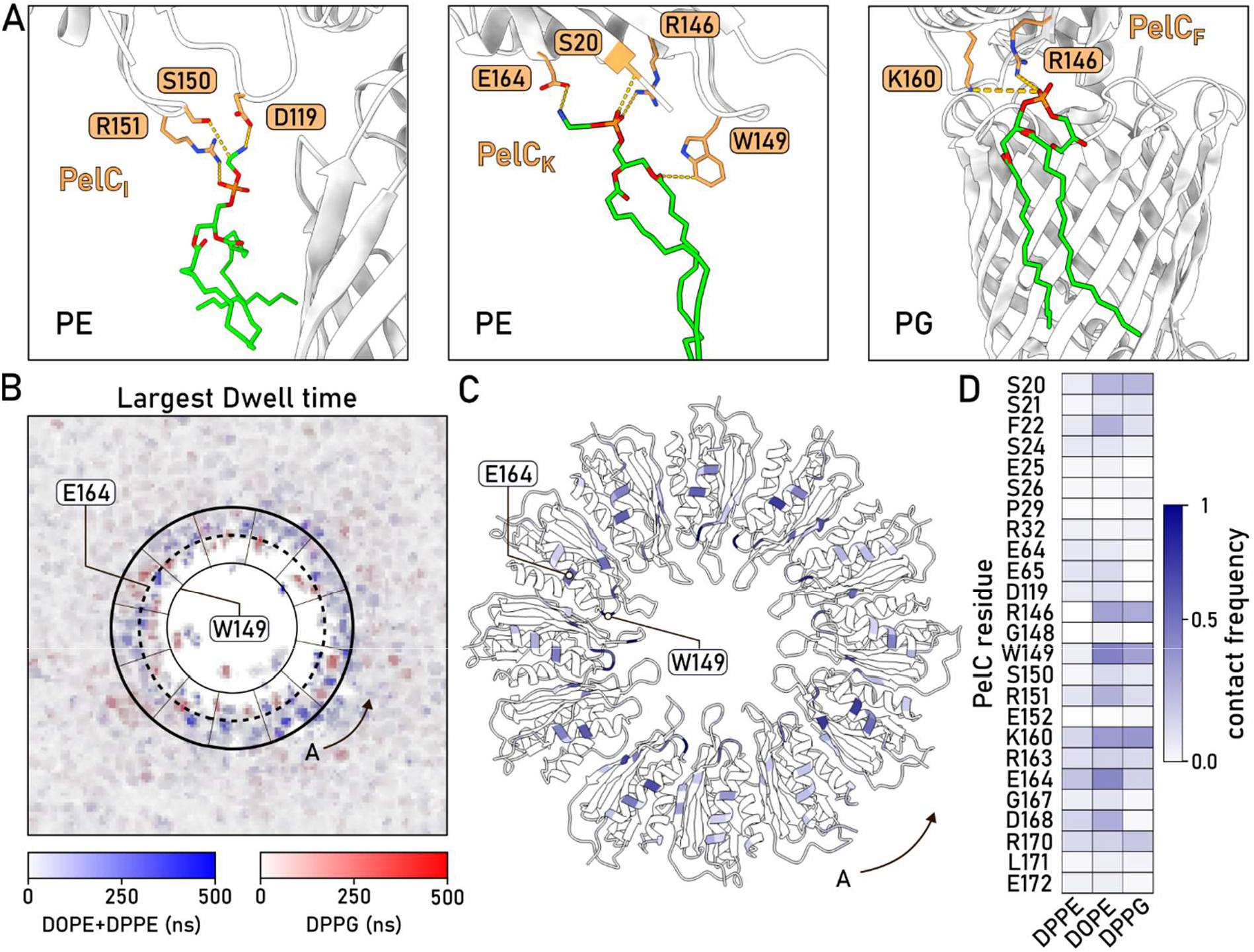
Contacts between the PelC ring and the periplasmic leaflet affect lipid dynamics. **(A)** Selection of lipid contacts (DOPE and DPPG) observed during MD simulations. **(B)** Plot for the largest dwell times of DOPE/DPPE and DPPG lipids during the MD simulation. The outer edge of the PelC ring is indicated as a solid bold line; the levels of the residues Trp-149 and Glu-164 are indicated as thin and dashed lines, respectively. Position of the PelC subunit A is indicated. **(C)** Distribution of the lipid contacts among the residues within the PelC ring (exemplified by the sumulation run #1). The residues are color-coded according to the frequency to form contacts with lipids. **(D)** Probability map of lipid-specific contact for PelC residues, averaged over all 12 subunits (exemplified by the sumulation run #1). 25 residues with the highest contact probability are shown.

## Conclusions

Assembly and dynamics of functional complexes in biological membranes rely on intricate protein:protein and protein:lipid interactions. Together with advances in structural biology, various membrane mimetics allow now to study how those interactions contribute to functional macromolecular architectures within the lipid bilayers. A particular attention is paid to the outer membrane of Gram-negative bacteria, as this asymmetric cell boundary contributes to the resistance against antimicrobial agents, and the embedded proteins facilitate numerous functions, from transport to adhesion, often associated with bacterial pathogenicity [28]. Here, we employed cryo-EM and all-atom MD simulations to examine the architecture and the interaction network of PelBC, the export machinery for Pel exopolysaccharide required for biofilm assembly in *P. aeruginosa*. We resolved the structure of the outer membrane complex after capturing it in two orthogonal systems, i.e. the large MSP-based nanodiscs reconstituted from detergent micelles and the polymer-based particles formed upon detergent-free extraction. Despite differences in the preparations, the reconstructions consistently showed the β-barrel of PelB associated with the dodecameric ring of the membrane-anchored PelC subunits. Improved local resolution in the MSP-based nanodiscs allowed modelling of the N-terminal fragment of PelB aligned with the gated exit site on the opposite side of the barrel. A chain of anionic residues spanning the barrel likely builds a route for the translocated polysaccharide. When extracted with cycloalkane-modified polymers, PelBC carried several lipid molecules from both membrane leaflets, making it possible to resolve lipid A molecule bound to the barrel, thus complementing the insights from synthetic lipid environment of MSP-based nanodiscs. To summarize, those results validate the earlier model of the complex, but also present the first application of cycloalkane-based polymers to study assemblies of outer membrane proteins, together with the associated lipids.

MD simulations of PelBC in the asymmetric bilayer that mimics the outer membrane of *P. aeruginosa* provided a complementary view on protein:lipid interactions. In agreement with the earlier analysis of PelB alone and the cryo-EM analysis of PelBC in UApol18, we observed lipid A molecules associated with the β-barrel via electrostatic interactions. With the PelC ring included in the simulations, we unveiled extensive protein:lipid interactions at the periplasmic side of the membrane. On the one hand, the simulations recapitulated positions of the lipid molecules at the PelB:PelC interface observed in cryo-EM reconstructions. The residence time of the docked lipids was dependent on the type of head group, demonstrating that electrostatic and steric factors at the interface confer interaction specificity. On the other hand, multiple PelC:lipid contacts were found at the periphery of the ring, matching the periodic density pattern observed in cryo-EM maps. The contacts were largely mediated by the charged residues of the C-terminal α-helix of PelC. The helix is essential for the functionality of the whole machinery, as its deletion abolishes the polysaccharide secretion without interfering with PelC expression [11]. While the earlier model suggested that the helix contributed to the translocation channel, such a role is ruled out by the resolved structures of PelBC. Instead, the helix mediates interactions with the proximate lipid membrane, which may be a prerequisite for the correct assembly and positioning of the PelBC complex.

Our study provides a comprehensive view on the unique architecture of the polysaccharide export machinery PelBC associated with microbial pathogenicity and the network of interactions it forms with the surrounding membrane. The results gained via the computational simulations corroborate the findings from cryo-EM analysis, and the combination may be further employed to decipher the route of the polysaccharide across the membrane. In its turn, the feasibility to extract the multi-subunit complex in a detergent-free manner opens new perspectives for studies, first of all for isolation from native membranes of *P. aeruginosa*. Such samples can serve not only for structural analysis, but also for lipidomics examination, so to identify the naturally occurring protein:lipid contacts formed along folding and assembly of the multi-subunit complex.

## Data availability

Structural data generated in this study have been deposited at the Electron Microscopy Data Base (EMDB) under accession numbers EMDB-58355 and Protein Data Bank (PDB) 31AF for PelBC-UApol18, and EMDB-58356 and PDB 31FB for PelBC-MSP2N2.

## Acknowledgements

We thank Susanne Rieder and Charlotte Ungewickell for technical assistance with the cryo-EM experiments, Melina Korbmacher and Thuy Linh Bui for contributions to sample preparation, and Jennifer Loschwitz for discussions. The research was supported by German Research Foundation (Deutsche Forschungsgemeinschaft, DFG), via the Research grant KE1879/6 and “Major Research Instrumentation” grant INST 208/939-1 FUGG (A.K.) and 510674444 (R.B). and “ACCeSS” project funded by the Ministry of Culture and Science of the State North Rhine-Westphalia (A.K.). J.R. and B.S. gratefully acknowledge the Gauss Centre for Supercomputing e.V. (www.gauss-centre.eu) for funding this project (pn98zo) by providing computing time on the GCS Supercomputer SuperMUC-NG at Leibniz Supercomputing Centre (www.lrz.de). Further computing time was granted by the Centre for Information and Media Technology at HHU Düsseldorf. We would like to acknowledge the Center for Advanced Imaging (CAi) at HHU Düsseldorf and especially Dr. Miriam Bäumers for supporting the negative-stain electron microscopy experiments.

## Author contributions

**CRH**: Investigation, Methodology, Visualization, Writing – original draft, Writing – review and editing

**MB**: Investigation, Methodology, Visualization, Writing – original draft, Writing – review and editing

**JR**: Investigation, Methodology, Visualization, Writing – original draft, Writing – review and editing

**OB**: Investigation, Data curation

**RB**: Funding acquisition, Project administration, Supervision, Writing – review and editing

**BS**: Conceptualization, Project administration, Supervision, Writing – review and editing

**AK**: Conceptualization, Funding acquisition, Project administration, Supervision, Writing – original draft, Writing – review and editing

## Methods

### Molecular cloning and protein expression

The pETDuet-1-based plasmid harbouring gene sequences encoding for *P. aeruginosa* PAO1 *pelB* (PA3063, residues 762-1193, including the pectate lyase signal peptide and an octa-histidine tag) and *pelC* (PA3062, including the signal peptide of *E. coli* LPP) was previously described [10]. Expression of the PelBC complex in *E. coli* and the crude membrane isolation were performed following the previously established protocol, without any modifications [10].

### PelBC isolation and reconstitution into nanodisc

The crude membrane extract containing the overexpressed PelBC was solubilized in 2% DDM (Glycon Biochemicals GmbH) in 50 mM HEPES pH 7.4, 300 mM KCl, 5% glycerol and adjusted to 10-fold volume relative to the membranes. After 1 h incubation, the non-solubilized material was removed by centrifugation at 21,000 x g for 15 min at 4 °C. The soluble fraction, supplemented with 5 mM imidazole, was loaded onto gravity-flow column packed with 450 µL pre-equilibrated Ni^2+^-NTA agarose beads. Upon incubation on the beads, for 1 h at 4 °C with rotating, the flow-through fraction was collected, and the resin was washed five times with 2 mL of buffer containing 40 mM imidazole and 0.05% DDM. The target complex was eluted in three fractions, each 800 µL, with IMAC buffer containing 300 mM imidazole and 0.05 % DDM. The pooled elution fractions were concentrated to 500 µL using Amicon Ultra-4 centrifugal filters, molecular weight cut-off 30 kDa (Merck/Millipore) prior injection onto Superdex 200 Increase 10/300 GL column connected to ÄKTA go system (Cytiva). The size exclusion chromatography (SEC) was performed in 50 mM HEPES pH 7.4, 150 mM NaCl, 0.03% DDM. For the mass photometry sample in LMNG, the SEC was performed in 50 mM HEPES pH 7.4, 150 mM NaCl, 0.003% LMNG. The protein concentration was determined spectrophotometrically based on calculated extinction coefficients of 497,430 M^-1^ x cm^-1^ for PelBC assuming 1:12 stoichiometry.

The nanodisc-forming scaffold protein MSP2N2 was expressed and isolated as described [13]. A lipid mixture composed of POPC:POPG lipids (molar ratio 70:30, Avanti Polar Lipids) was prepared in chloroform, the solvent was evaporated, and the lipids were suspended to the final concentration of 5 mM in 50 mM HEPES pH 7.4 and 150 mM NaCl. The formed liposomes were extruded stepwise through 1 µm and 200 nm track-etch membranes (Whatman/Cytiva) and 0.5% DDM was added to solubilize the lipids upon incubation at 40 °C for 15 min. Subsequently, the purified PelBC complex was mixed with the scaffold protein MSP2N2 and the lipids at the molar ratio of 1:4:500 followed by 20 min incubation on ice. After the incubation, pre-washed Bio-Beads SM-2 sorbent (Bio-Rad Laboratories) was added and incubated overnight at 4 °C upon rotation. The reconstitution reaction was loaded onto Superose 6 Increase 10/300 GL column connected to ÄKTA pure system (Cytiva), and SEC was performed in 50 mM HEPES pH 7.4, 150 mM NaCl. The elution fractions were analyzed via SDS-PAGE. The fractions containing the assembled PelBC complex in nanodisc were pooled, concentrated, and stored flash-frozen for cryo-EM analysis.

### Polymer-based isolation of PelBC

The crude membrane extract containing overexpressed PelBC was solubilized using 2.5% UApol18 (Cube Biotech GmbH) in 50 mM HEPES pH 7.4, 300 mM KCl, 5% glycerol, and adjusted to 4-fold volume relative to the membranes. After incubation overnight at 4 °C with rotation, the non-soluble material was removed by centrifugation at 21,000 x g for 15 min at 4 °C. The soluble fraction was supplemented with 2 mM histidine and loaded onto a gravity-flow column packed with 400 µL pre-equilibrated Ni^2+^-NTA agarose beads (Protino, Macherey-Nagel). Binding was performed for 1 h 15 min at 4 °C upon rotation. After removing the flow-through fraction, the resin was washed five times with 2 mL buffer containing 20 mM histidine without additional UApol18. The target material was eluted with the buffer containing 200 mM histidine in three fractions, each of 800 µL. To reduce the histidine concentration prior cryo-EM analysis, the pooled fractions were concentrated three times to 100 µL and diluted back to 4 mL with 50 mM HEPES pH 7.4, 150 mM NaCl. The protein concentration was determined spectrophotometrically as described before. After the final concentration step, the sample was flash-frozen and stored for mass photometry and cryo-EM analysis. To evaluate the homogeneity and integrity of the PelBC complex by negative-stain EM, a 3 µL drop (0.1 mg/ml protein concentration) was applied onto a copper grid (Carbon Support Film 200 Mesh, CF200-CU-50, Electron Microscopy Sciences) and left to soak in for 1 min. The liquid was carefully removed, and the grid was placed on a drop of uranyl acetate for 1 min for staining. The grid was then left to air-dry for at least 20 min prior loaded into a Zeiss EM902 transmission electron microscope. Mass photometry measurements were carried out using Two^MP^ instrument (Refeyn Ltd.). Calibration in the mass range from 90 kDa to 1 MDa was performed using MassFerence™ P1 kit (Refeyn Ltd.). Flash-frozen aliquots of the samples were thawed and thermally equilibrated for 10 minutes before measurement. The optimal protein concentration was determined experimentally by successive dilution with the buffer without LMNG. 10 µL of the sample was loaded in gasket wells and measured using an acquisition time of 60 s. The results were plotted as histograms, and the molecular weight determination was performed using the instrument’s software.

### Cryo-EM sample preparation and data collection

An aliquot of 3.5 µL of PelBC reconstituted in MSP2N2 or UApol18 nanodiscs was applied to glow-discharged Quantifoil Cu 300 mesh R2/1 (MSP2N2) or R3/3 (UApol18) grids with an additional 2 nm layer of carbon, after being supplemented with (1H, 1H, 2H, 2H-perfluorooctyl)-β-D-maltopyranoside (Anatrace) to a final concentration of 0.03 %. Following a waiting time of 45 s, the grids were blotted for 3 s and plunge frozen in liquid ethane using a Vitrobot Mark IV (Thermo Fisher). Data collection was performed at 300 keV using a Titan Krios microscope equipped with a Falcon 4i direct electron detector and a SelectrisX energy filter (all Thermo Fisher), at a pixel size of 0.727 Â. Dose-fractioned movies were collected in a defocus range from −0.5 to 3.0 um and with a total dose of 40 e-per Â^2^, fractionated in 40 frames to obtain a total dose of 1 e^-^ per A^2^ per frame.

### Cryo-EM data processing

Gain correction, movie alignment and summation of movie frames was performed using MotionCor2(ref). Further processing was carried out in cryoSPARC v4.6 [29]. For PelBC in MSP2N2-based nanodiscs, the CTF parameters of the full data set consisting of 26,999 micrographs, were estimated using PatchCTF [30]. Using Blob Picker and 2D classification, 82,455 particles were picked from a subset of around 13,000 micrographs and later used to train a TOPAZ [31] model for particle picking on the full data set. A total of 3,028,720 particles were picked and cleaned through successive rounds of 2D classification. A subset of 737,113 particles, belonging to the best resolved classes, was extracted, binned twice and used as input for an *ab initio* reconstruction job with 3 classes. Multiple rounds of heterogeneous refinement led to a well resolved class of 476,522 particles, later refined in a Homogeneous Refinement job. A mask covering the β-barrel of PelB was used to perform focused classification into 3 classes. From the resulting reconstructions, one class of 194,769 particles displayed well resolved details on the PelC ring and PelB β-barrel. These particles were extracted at 0.727 A/px and by setting the filter resolution parameter at 3.0 Å, focused 3D classification allowed further sorting into 3 classes. The best resolved class, consisting of 107,242 particles, was refined to final resolution of 2.76 Å.

For PelBC in UApol18, a total of 12,644 micrographs were CTF corrected using PatchCTF. A set of 1,539,647 particles was picked on 6,585 micrographs using Blob Picker and cleaned by 2D classification. 407,787 particles belonging to well-resolved 2D classes were used to train a TOPAZ model which, after picking on the full data set, resulted in 730,891 particles. After successive rounds of 2D classification, a cleaned set of 596,463 was used to generate a consensus refinement using a Homogeneous Refinement job. Focused classification with a mask around the β-barrel of PelB resulted in four classes displaying a well-resolved PelC ring, but different levels of map quality for the β-barrel. The best-resolved classes were combined and used to generate a new consensus refinement, composed of 309,144 particles. Further sorting into 3 classes was achieved by performing a focused 3D classification, setting the filter resolution at 4 Å. The class with the best β-barrel and ring features, containing 119,846 particles, was finally re-extracted at 0.727 A/px and refined to final resolution of 2.61 Å.

### Model building

The PelBC complex cryo-EM structure in MSP1D1 nanodisc (PDB ID 9H80; [10]) was used as initial model and fitted into the cryo-EM volumes. Manual adjustments were performed using the COOT program (WinCoot version 0.9.8.92, [32]) and the ISOLDE plugin implemented in ChimeraX (version 1.10.1, [33]), with ongoing refinement achieved through the option real space refinement in the PHENIX program (version 1.20.1-4487, [34]). To model the PelC anchor chains, a modified cysteine was modelled using the Ligand Reader & Modeler of CHARMM-GUI [35, 36]. The length of each anchor was modelled for individual PelC subunits based on the present cryo-EM volume. The ligand coordinates acquired from PDBs were prepared using the option elbow in the PHENIX program [37], also to receive the cif files for refinement. The atom names of the cysteine in the PDB and cif files were changed afterwards manually to match the names of a normal cysteine (C1 to Cα, C2 to C, etc.). Additionally, the cif-classification was changed from a ligand to a modified peptide. Next, the Cys-19 in the structure model were replaced with the modified cysteines and for each subunit placed within the cryo-EM volume. The modified cysteines were parametrized based on ANTECHAMBER implemented in the ISOLDE plugin, fitted into the cryo-EM densities for each subunit and connected to the protein model with bond restrains after real-space refinement in PHENIX. As definitive evidence cannot be provided as to whether these lipids are co-purified from the *E. coli* outer membrane or originate from the nanodisc reconstitution process, phosphatidylethanolamine (PE) molecules were modelled into regions where the density was of sufficient quality.

### Molecular dynamics simulations

The entire simulation model was constructed using the Membrane Builder module of the CHARMM-GUI webserver [35, 38–40]. The MSP1D1 nanodisc structure of the PelBC complex was used as the basis for the simulation model [10]. The PelC subunits were modified with a lipid tail post-translational modification on residue Cys-19 to CYSL (triacylation). The modification was selected as it most closely resembles the density observed in the EM structure. For the membrane, a heterogeneous bilayer was chosen to simulate the outer membrane composition of *P. aeruginosa*. The lipid leaflet facing the extracellular space consists entirely of LPS molecules, which were generated using the LPS Modeler of CHARMM-GUI with the Type 1 lipid for *P. aeruginosa* [35, 39]. The leaflet facing the periplasmic space is composed of a mixture of DOPE, DPPG, and DPPE lipids in a ratio of 46:31:23 [27]. The PelBC protein complex was initially positioned 8.5 Å above the membrane to allow it to settle more naturally, enabling the surrounding lipids to accommodate the membrane protein.

To neutralize the LPS charges, Ca^2+^ ions were added to the system. Conditions mimicking the natural environment of the outer membrane of *P. aeruginosa* were applied, with a NaCl concentration of 25 mM, a temperature of 298.15 K, and a pressure of 1 atm. To investigate the lipid density beneath chain I of the PelC subunits, two systems were prepared, one with a docked DPPE lipid molecule (simulated in triplicates) and one with a docked DPPG lipid placed at the observed density site. Docking of the lipids to the protein structure was carried out using AutoDock Vina (version 1.2.3, [41, 42]), and the resulting coordinates were subsequently used to reposition the corresponding lipid within the simulation model. The resulting simulation box had dimensions of 18.59 nm, 18.59 nm, 16.43 nm (x, y, z) and contained 504,179 atoms.

For the MD simulations, GROMACS 2024.3 [43, 44] was used with the Charmm36m force field for the proteins [45, 46], the Charmm36 lipid force field [47], and the Charmm-modified TIP3P water model [48]. Prior to the production run, energy minimization and multistep equilibration were performed to relax the system. Using the steepest descent algorithm, the system’s energy was reduced to 1000 kJ mol^−1^ nm^−1^ during energy minimization. The system was then equilibrated in six sequential steps: the first two steps (125 ps each, 1 fs time step) were run in the NVT ensemble with a Berendsen thermostat (298.15 K) [49], followed by four NPT steps (125, 500, 500 and 500 ps; time step increased to 2 fs from the third NPT step onward) using a V-rescale thermostat (298.15 K) [50] and semi-isotropic C-rescale pressure coupling [51]. Harmonic position restraints on the protein backbone, side chains, and lipid head groups, together with dihedral restraints on the protein, were applied during minimization and progressively released over the course of equilibration. All bonds involving hydrogen atoms were constrained with the LINCS method [52]. Electrostatic interactions were treated using the particle-mesh Ewald (PME) method [53] with a real-space cutoff of 1.2 nm, and van der Waals interactions were handled using a force-switch function between 1.0 and 1.2 nm. For the production run, the Nosé-Hoover thermostat [54] and the semi-isotropic Parrinello-Rahman barostat [55] were employed. The integration time step was set to 2 fs, and frames were saved every 20 ps. In total, three 500 ns simulations were performed with a docked DPPE molecule positioned near chain I, and one 350 ns simulation was conducted with a docked DPPG lipid (instead of DPPE) placed at the same site.

To assess the stability of the protein complex during the simulation, RMSD and RMSF were calculated using the GROMACS built-in functions *gmx rms* and *gmx rmsf*. Lipid residence times were analyzed using the software suite MOSAIC (v1.10, [56]), in which the positions of the lipid head groups were tracked over the entire simulation time of 500 ns or 350 ns with a stride of 1 ns. Here, the grid spacing was set to 0.02 nm) and the cutoff radius (*r*_cut_) to 0.3 nm. Contact frequencies between the lipids and the protein residues were determined using the GetContacts software [57], applying first the *get_dynamic_contacts*.*py* function, followed by *get_contact_frequencies*.*py*. All plots were generated with the Python libraries Matplotlib, Seaborn, and NumPy.

### Software

Protein structures were visualized, analyzed, and rendered for figures using ChimeraX (version 1.10.1, UCSF). Figures were assembled using Inkscape (v1.3.1).

