## Supplemental Figures 1-19 for "Exopolysaccharide export complex PelBC of *Pseudomonas aeruginosa* attenuates the dynamics of the surrounding outer membrane"

### Supplementary data

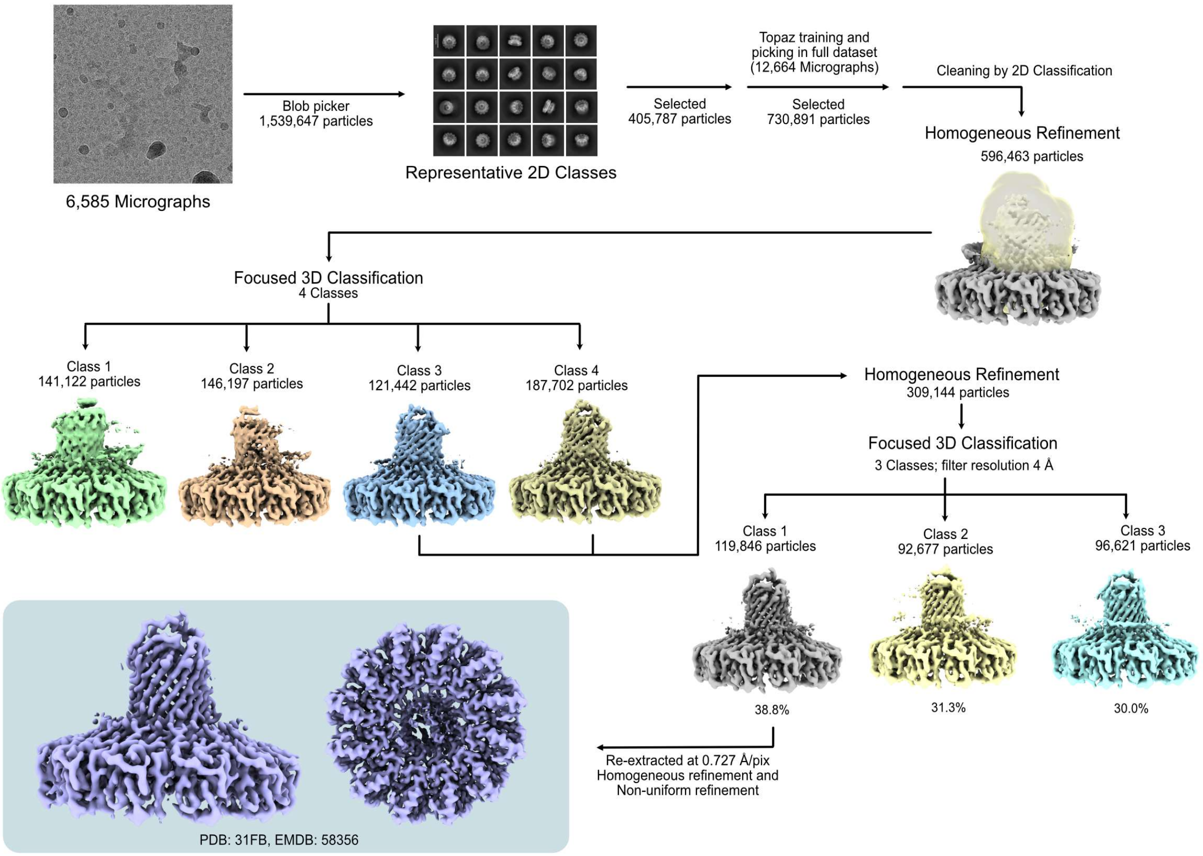


**Supplemental Figure 1. Single-particle analysis towards the PelBC structure in MSP2N2-based nanodiscs.** Summary of the sorting/refinement procedures performed upon single-particle analysis of the cryo-EM data set collected on the nanodisc-reconstituted PelBC complex.

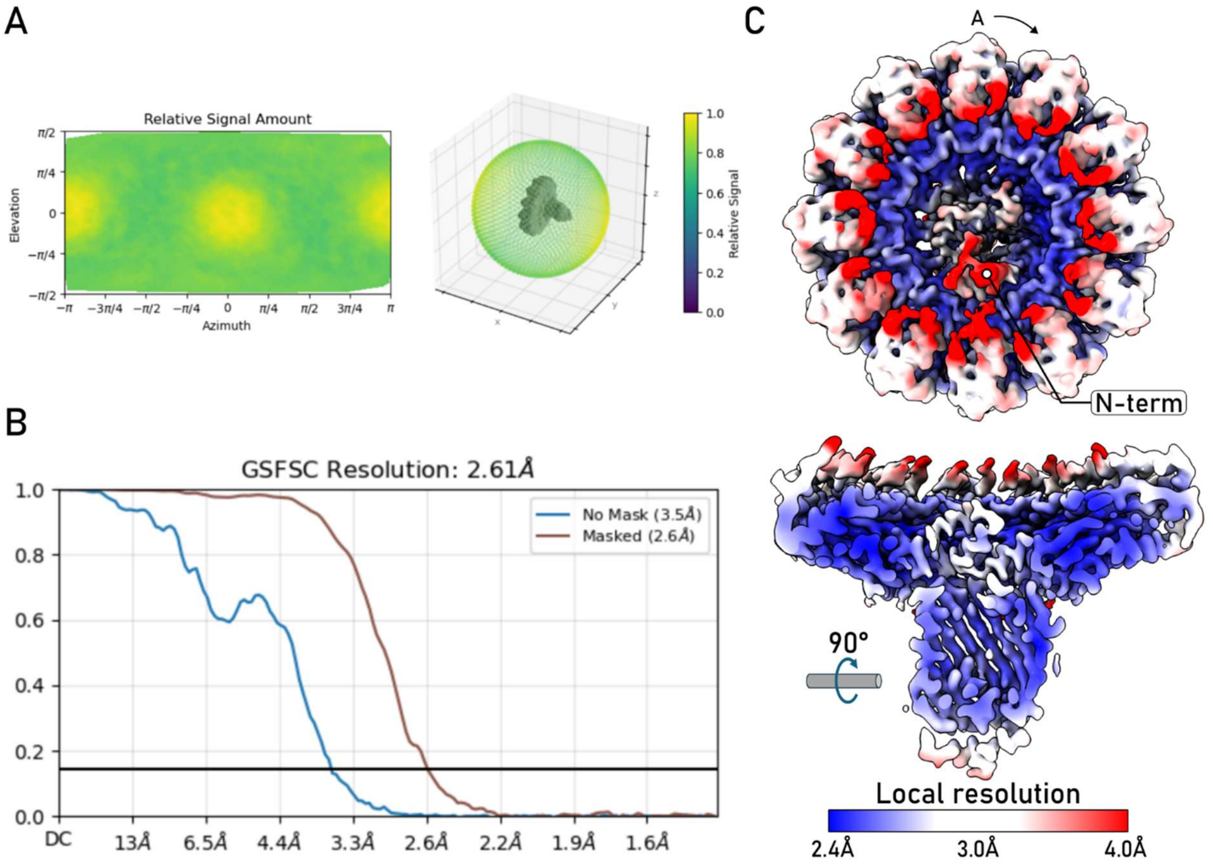


**Supplemental Figure 2. Global and local resolution of PelBC in MSP2N2-based nanodiscs.**

**(A)** Angular distribution plot for the final reconstruction obtained from cryoSPARC.

**(B)** Gold-standard Fourier shell correlation resolution curve for the final map, displaying resolution with a mask automatically generated by cryoSPARC.

**(C)** Final PelBC map colored according to the local resolution as determined by cryoSPARC. **Top:** View from the periplasmic side. PelC subunit A and the position of the N-terminal domain of PelB are indicated. **Bottom:** Cross-section of the complex, orthogonal to the membrane plane.

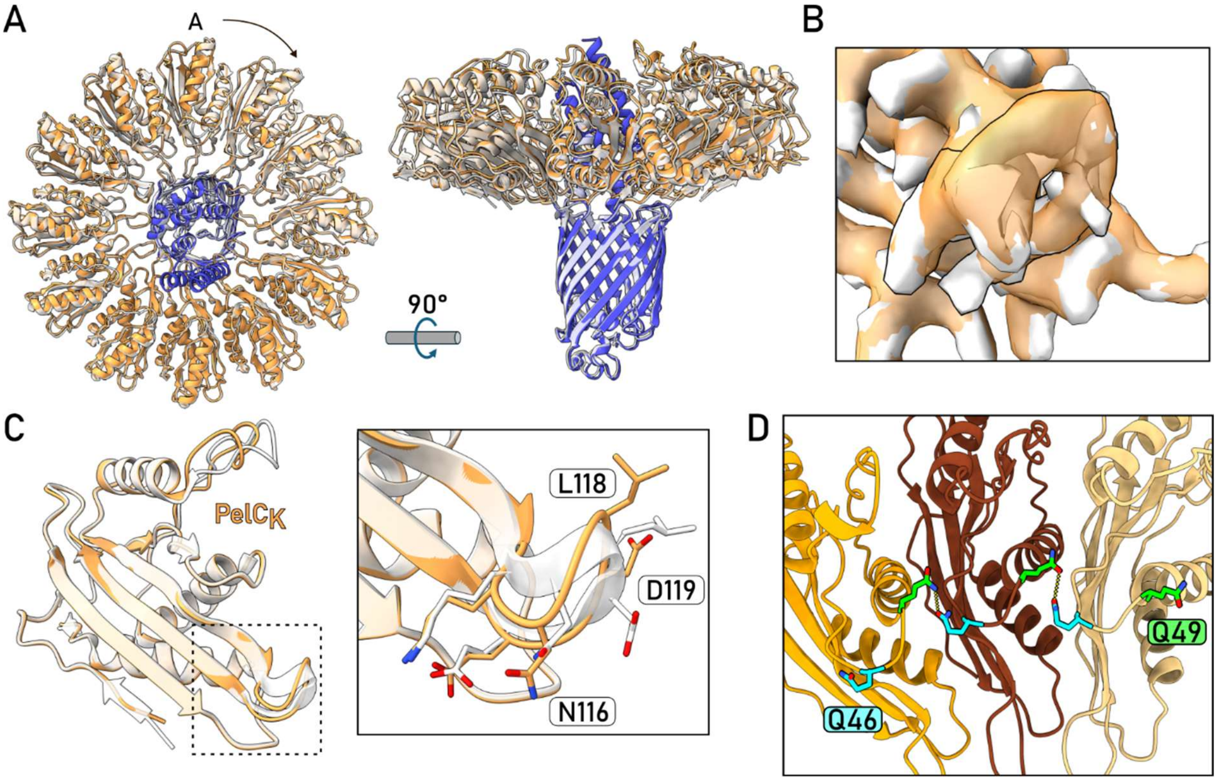


**Supplemental Figure 3. Structural model of PelBC in MSP2N2-based nanodiscs.**

**(A)** PelBC manifests nearly identical structures when reconstituted into MSP2N2-based lipid nanodiscs (PelB shown in blue, PelC in orange) and MSP1D1 (PDB ID 9H80; shown in white).

**(B)** Cryo-EM density of the PelC$_K$ D-loop in MSP2N2- and MSP1D1-based nanodiscs (shown in orange and white, respectively).

**(C)** Modelled conformations of PelC$_K$ in MSP2N2- and MSP1D1-based nanodiscs (shown in orange and white, respectively). D-loop is shown in magnification.

**(D)** Interaction of Gln-46 and Gln-49 between neighbouring PelC subunits

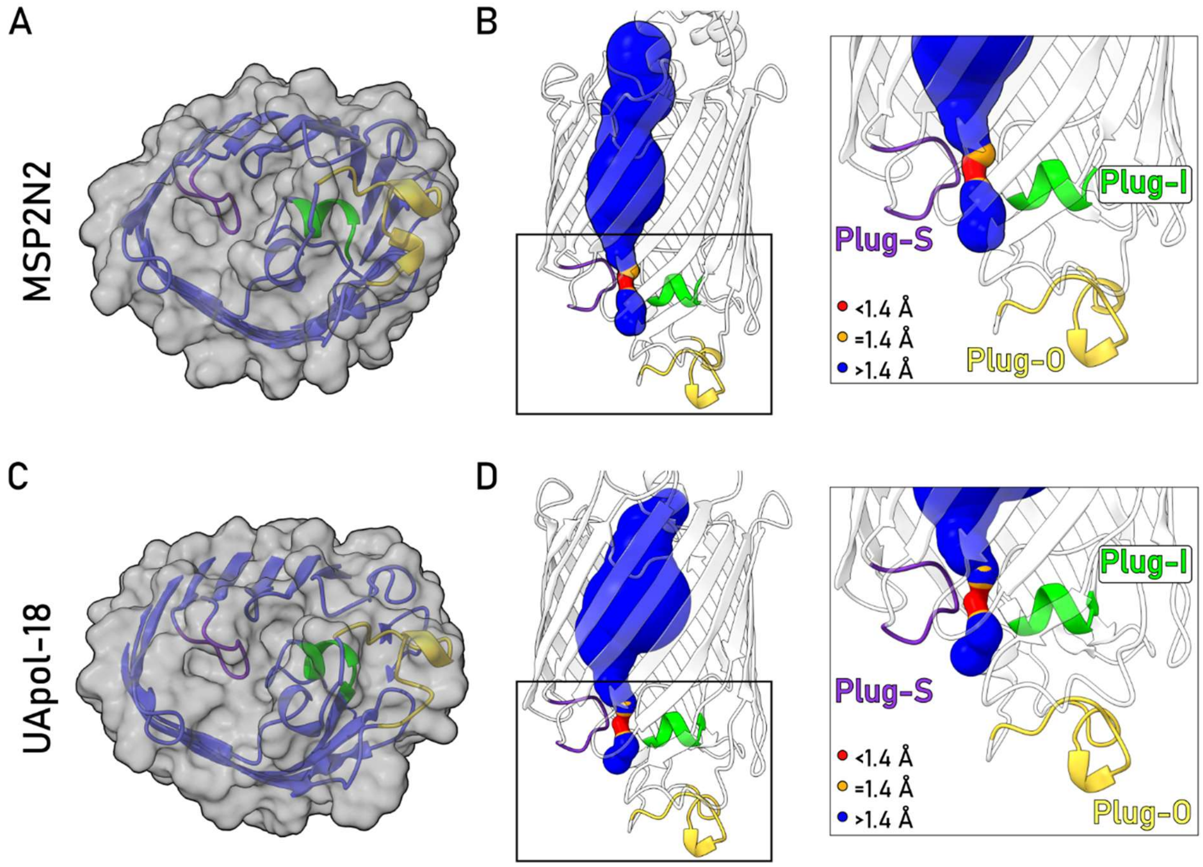


**Supplemental Figure 4. PelB manifests closed conformation of the exit site.**

**(A)** Surface representation of PelB in MSP2N2-based nanodiscs suggests a closed conformation of the β-barrel.

**(B)** The tunnel within the PelB β-barrel in MSP2N2-based nanodiscs color-coded according to its width. The bottleneck (red) does not allow passage of a water molecule (van der Waals radius 1.4 Å). The tunnel was modelled with MOLE 2.5 ([1], Pore mode: Beta structure on, membrane region off, interior threshold 1.4 Å, probe radius 5).

**(C and D)** Same as (A and B), but for PelBC isolated with UApol18.

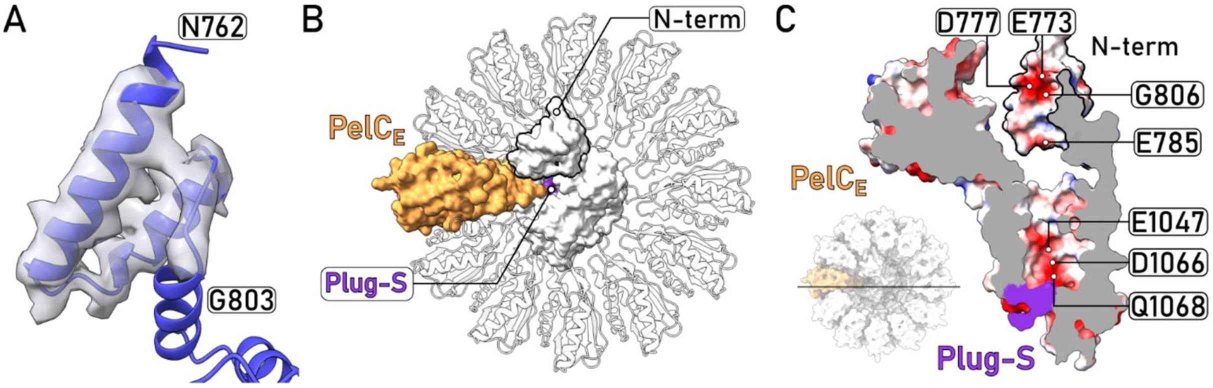


**Supplemental Figure 5. The N-terminal TPRL domain of PelB is aligned with the putative exit site.**

**(A)** The electron density corresponding to the N-terminal domain of PelB (residues 762-803) resolved in the MSP2N2-based nanodiscs.

**(B)** The N-terminal domain of PelB and PelC subunit E (PelC$_E$) form a consolidated groove aligned with the putative exit site at the extracellular side of the β-barrel (closed by the loop Plug-S in the resolved structures).

**(C)** Cross-section view of PelB and PelC$_E$ showing the electrostatic potential distribution. The anionic residues of the N-terminal domain and the β-barrel of PelB are aligned with Plug-S, possibly marking a route for the positively charged Pel exopolysaccharide.

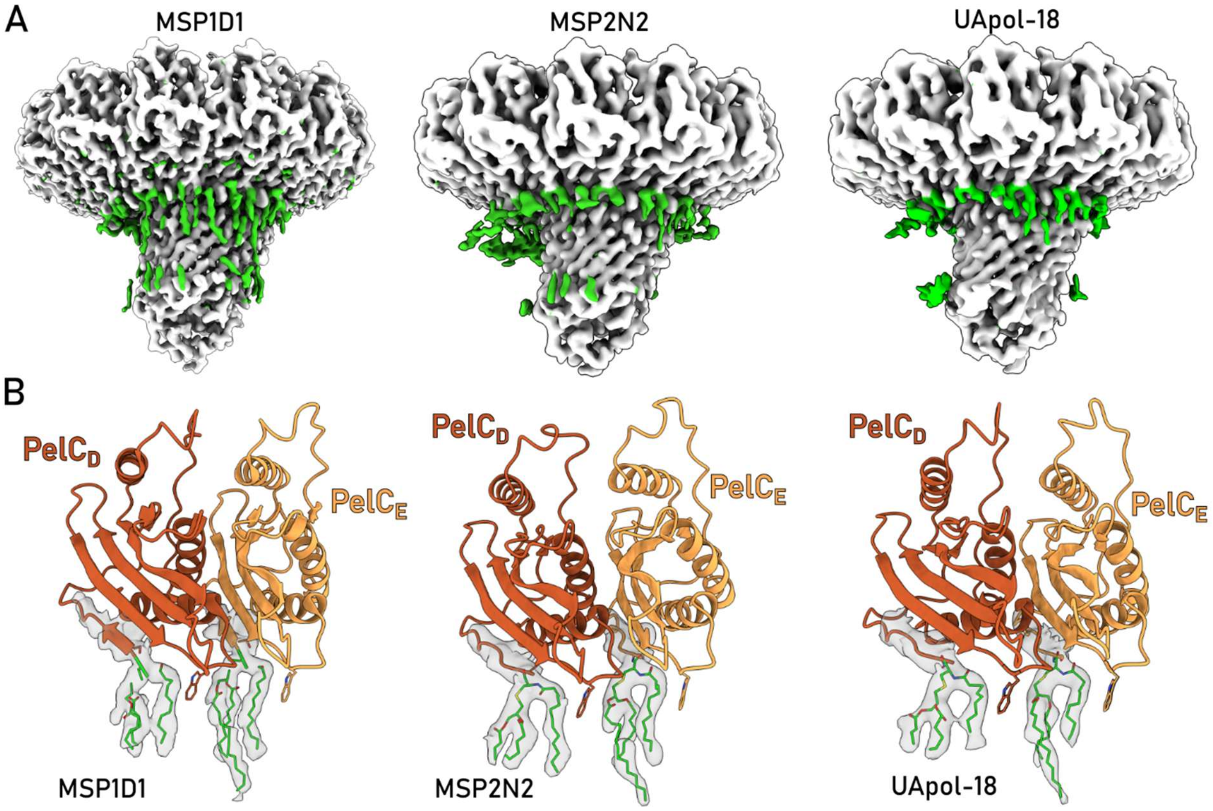


**Supplemental Figure 6. Lipid-like densities in cryo-EM reconstructions of PelBC.**

**(A)** Cryo-EM reconstructions of all three environments showing the extra densities (green) which correspond to the associated lipid molecules.

**(B)** Lipid anchors of PelC clamped between Tpr-149 in all three environments, shown on examples of PelC subunits D and E.

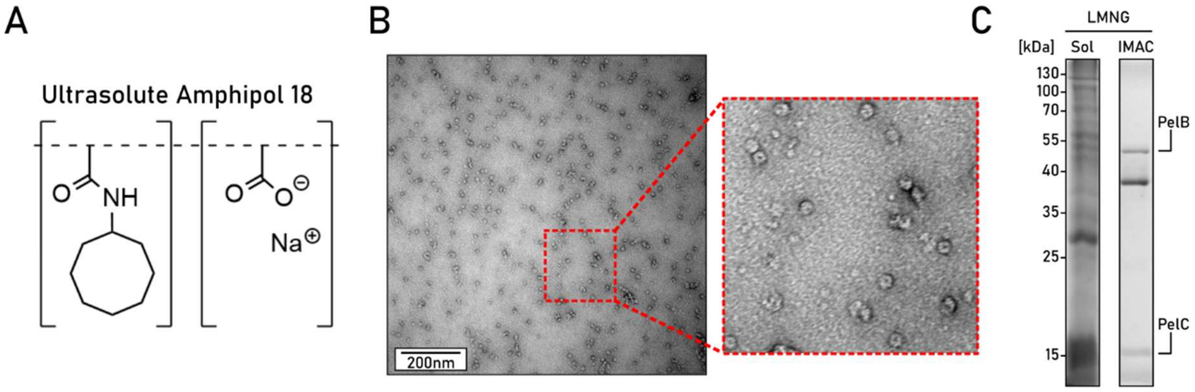


**Supplemental Figure 7. Cycloalkane-modified amphipol UApol18 allows isolation of the intact PelBC complex.**

**(A)** Chemical structure of the UApol18 polymer [2].

**(B)** A representative negative-stain EM micrograph of the PelBC-UApol18 sample after IMAC purification (captured at 85,000x magnification). Donut-shaped particles (zoomed view) indicate presence of the assembled PelC ring.

**(C)** SDS-PAGE of the PelBC complex solubilized (“Sol”) and isolated with LMNG as a reference for the mass photometry measurements.

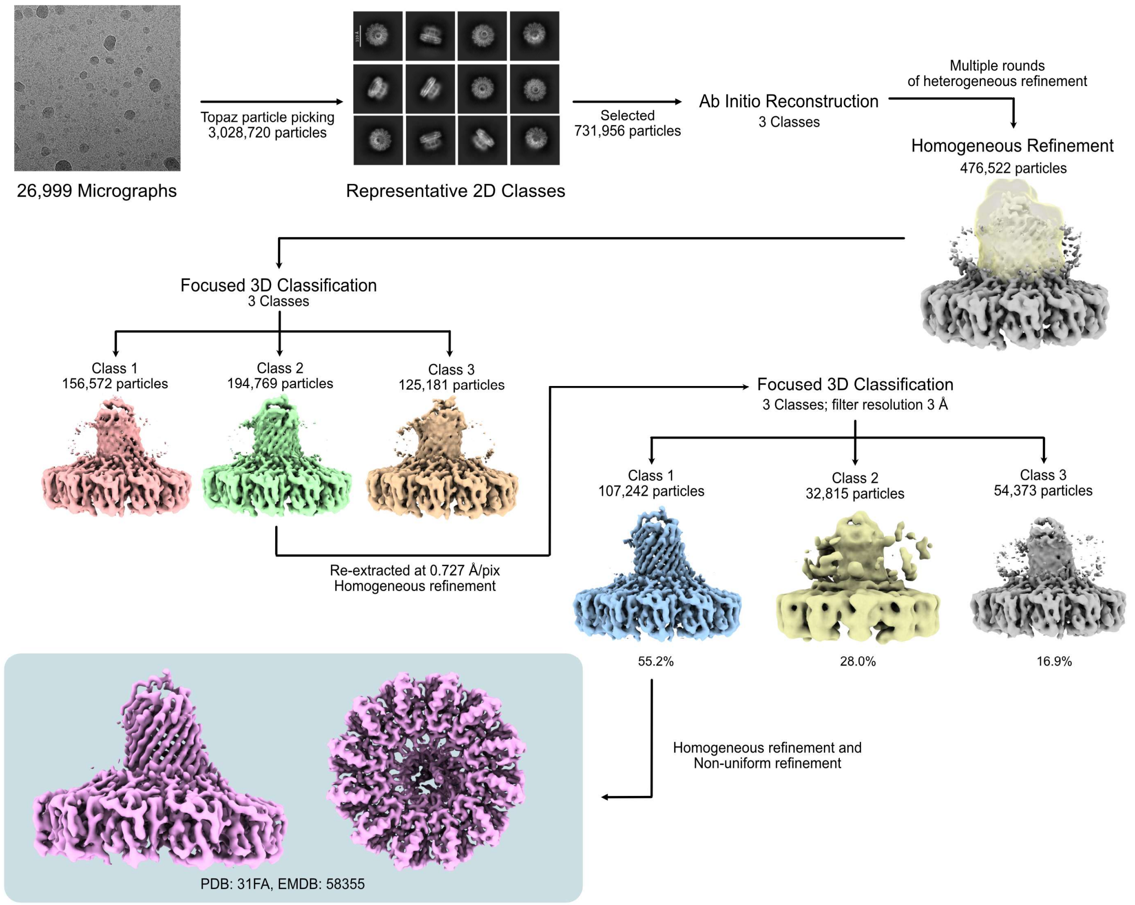


**Supplemental Figure 8. Single-particle analysis towards the PelBC structure in UApol18.** Summary of the sorting/refinement procedures performed upon single-particle analysis of the cryo-EM data set collected on the PelBC complex isolated using UApol18 polymer.

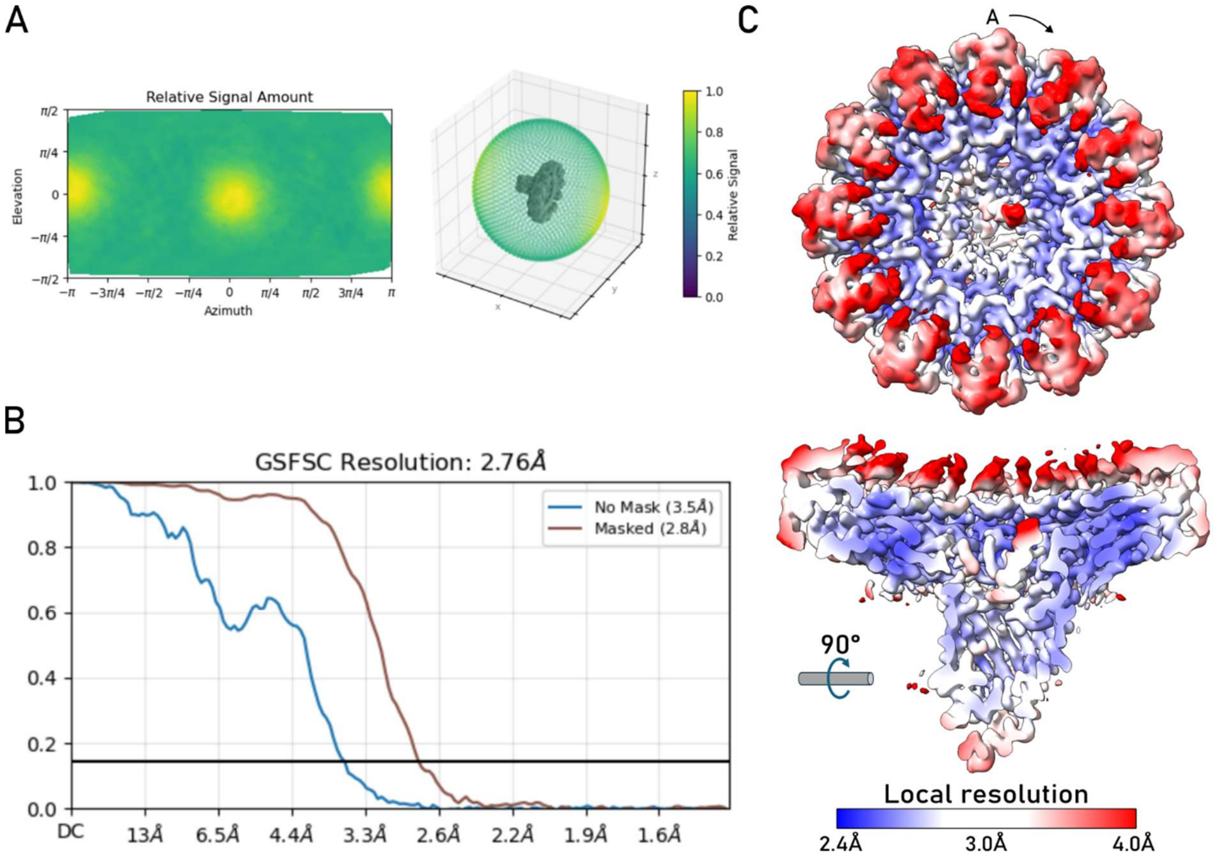


**Supplemental Figure 9. Global and local resolution of the PelBC-UApol18 map.**

**(A)** Angular distribution plot for the final reconstruction obtained from cryoSPARC.

**(B)** Gold-standard Fourier shell correlation resolution curve for the final map, displaying resolution with a mask automatically generated by cryoSPARC.

**(C)** Final PelBC map colored according to the local resolution as determined by cryoSPARC. **Top:** View from the periplasmic side. PelC subunit A is indicated. **Bottom:** Cross-section of the complex, orthogonal to the membrane plane.

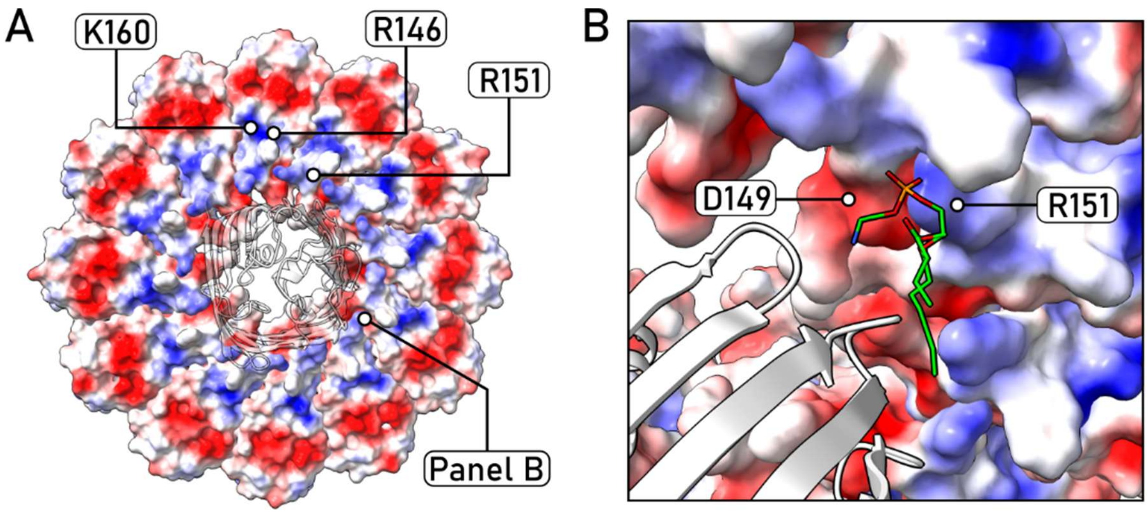


**Supplemental Figure 10. Electrostatic potential of the PelC ring.**

**(A)** View on the electrostatics potential of the PelC ring (membrane-facing side), with the residues R146, R151 and K160 highlighted.

**(B)** Cryo-EM resolves a PE lipid forming electrostatic interactions with PelC subunit I, with the amine group oriented to Asp-149 and the phosphate to Arg-151.

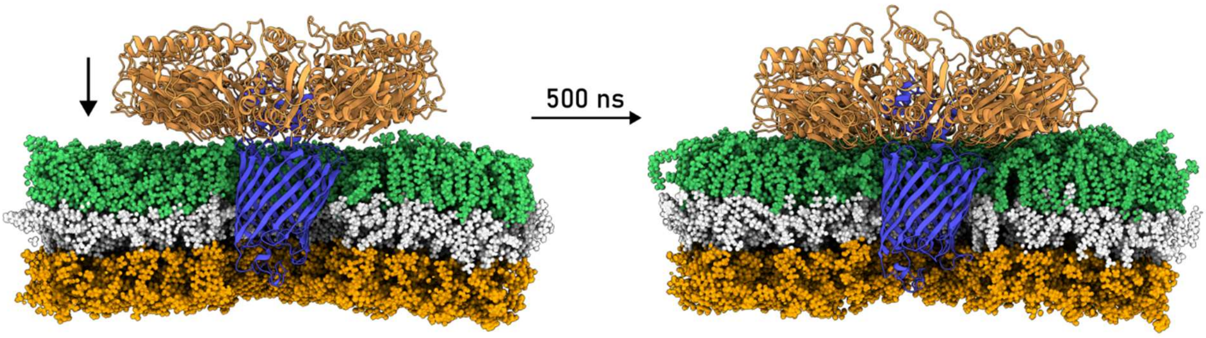


**Supplemental Figure 11. PelBC spontaneously accommodates itself within the modelled outer membrane of *P. aeruginosa*.**

**Left:** Position of PelBC at the start of the production run (after energy minimization and equilibration). **Right:** Position of PelBC at the end of the production run (duration of 500 ns).

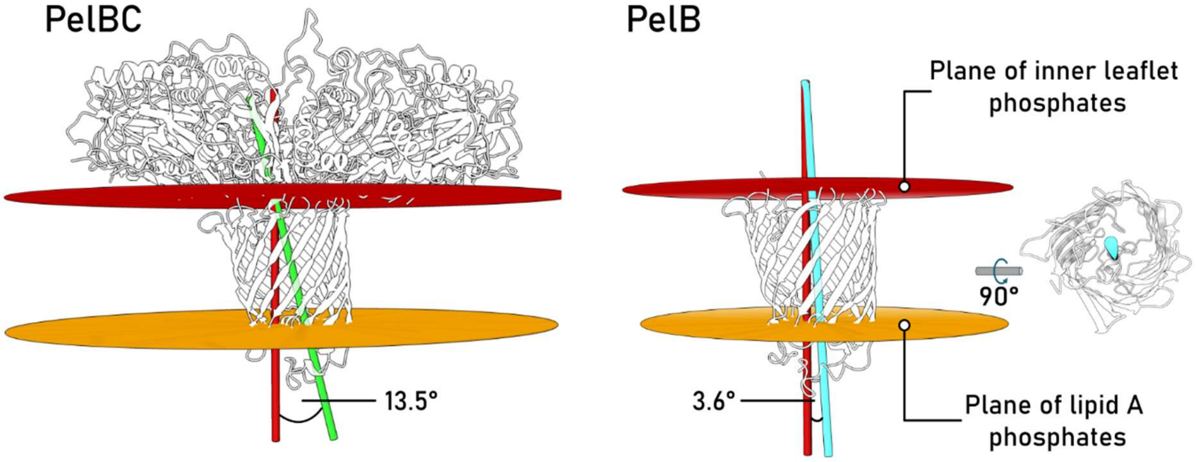


**Supplemental Figure 12. Orientation of the PelB ß-barrel in simulated membranes.**

The PelB β-barrel manifests different tilts within the membrane if simulated within the PelBC complex (left) or alone (right, [3]). The tilt measurements were performed using ChimeraX. Phosphates of the inner leaflet lipids (red) and phosphates of the lipid A molecules in the outer leaflet (orange) were selected to define planes of the inner and outer borders of the lipid bilayer. For the tilt of the barrel, two centroids were defined based on the residues at the inner leaflet and the outer leaflet plane to define a tilt axis (green and cyan) through these two centroids. The angle between the normal of the inner leaflet plane (red) and the tilt axis was then measured by ChimeraX.

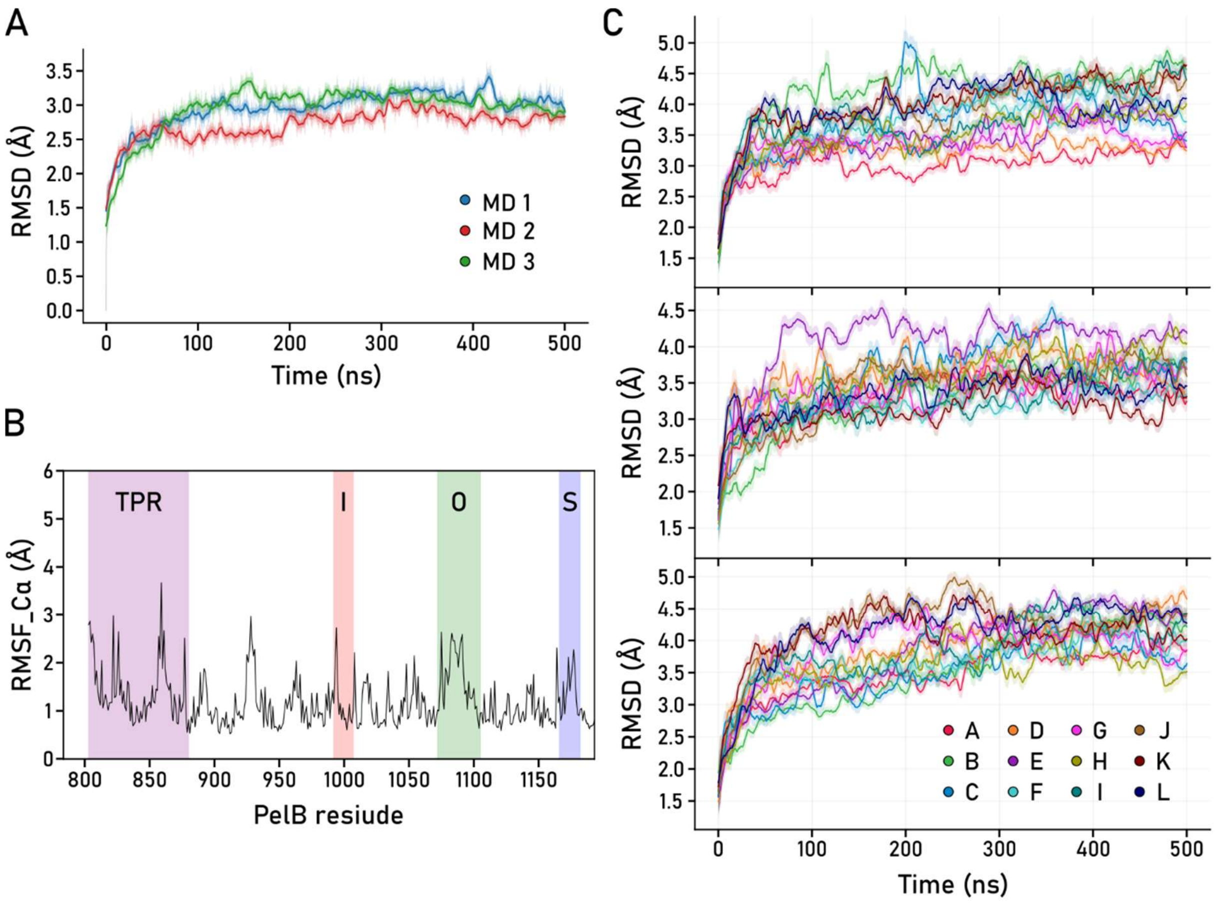


**Supplemental Figure 13. Conformational flexibility of PelBC subunits.**

**(A)** RMSD plot of PelB. The dark line is the 5 ns time mean and the faint colour shows the standard deviation. In all three simulations is no increased RMSD value noticeable, providing evidence that PelB protein is stable in the simulated membrane over the simulation time (500 ns).

**(B)** Mean RMSF plot of PelB over the three simulations. Increased flexibility can be identified in the TPR (purple), Plug-O (green) and Plug-S region (blue), as well as in the loop between β-strand 3 and 4 (residues 924-936).

**(C)** RMSD plots of the PelC subunits overlaid, all three simulations represented. The dark line is the 5 ns time mean and the faint color shows the standard deviation. No distinguished increase in RMSD values over time, showing that the PelC subunits are stable over the simulation time (500 ns).

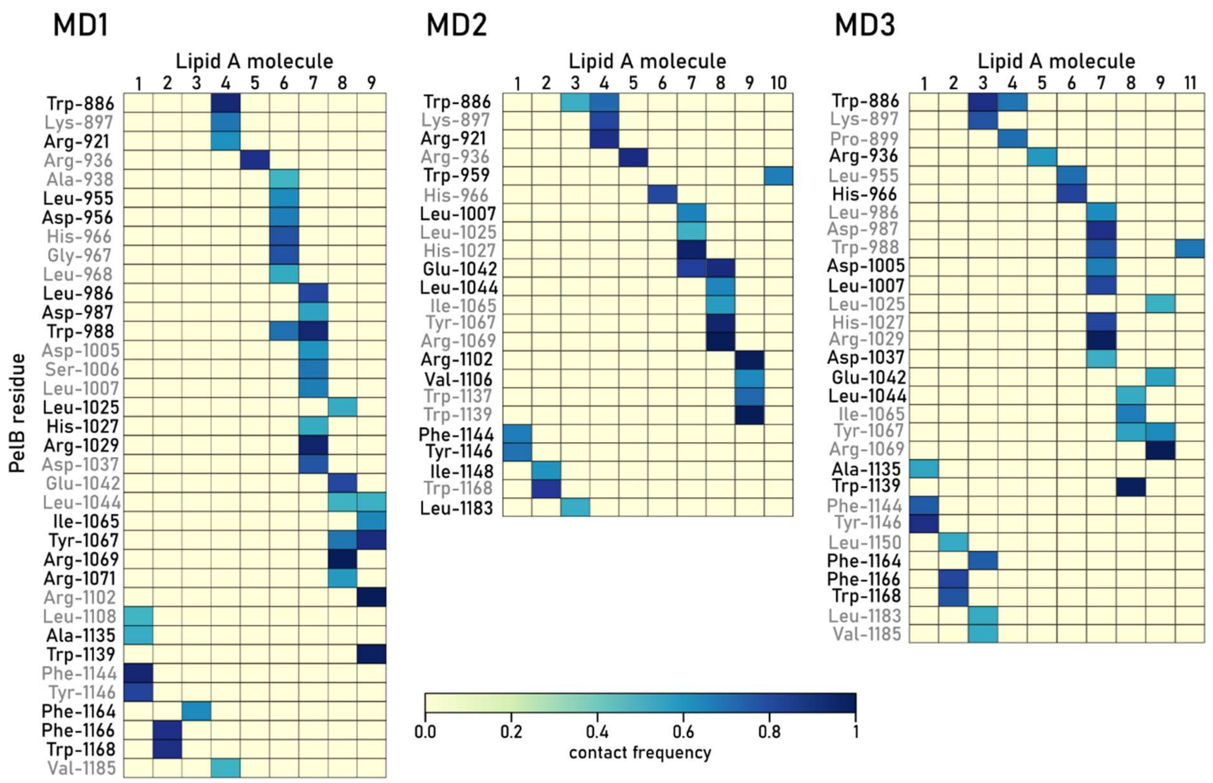


**Supplemental Figure 14. Contact frequency plot of the PelB residues interacting with lipid A molecules during the simulation time**.

Residues originating from the same β-strand are grouped by the label colours. Lipid A molecule 1 to 9 are numbered as shown in Figure 4, number 10 and 11 are only present in simulation 2 or 3 and would appear between molecule 5 and 6 (number 10) and between molecule 6 and 7 (number 11). As they are not located in direct proximity to the barrel, the possibility to interact with the PelB residues are relatively low.

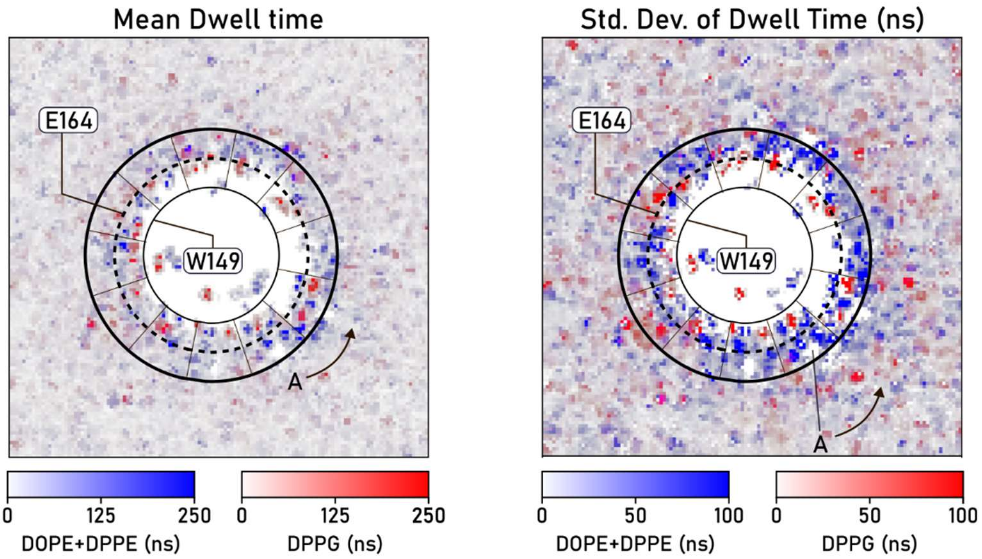


**Supplemental Figure 15. Lipid mobility under the PelC ring.**

Lipid residence time plots for the mean dwell time of simulation 1 and the corresponding standard deviation showing the localisation of PE and PG lipids. The outer edge of the PelC ring is indicated as a solid bold line; the levels of the residues Trp-149 and Glu-164 are indicated as thin and dashed lines, respectively. Position of the PelC subunit A is indicated.

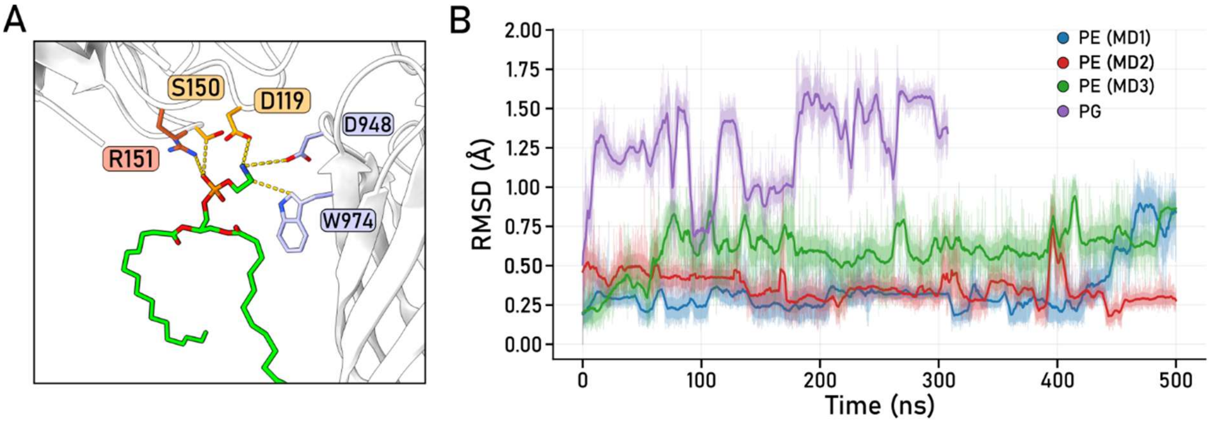


**Supplemental Figure 16. Headgroup-specific interactions of lipids with PelBC.**

**(A)** A docked DPPE lipid shown interacting with PelB residues and PelC subunit I (yellow) and H (orange).

**(B)** RMSD plot of the headgroups of docked PE and PG lipids under PelC subunit I and H. The plot suggests stable docking of PE lipids (in all three simulations) while the PG lipid is moving significantly more.

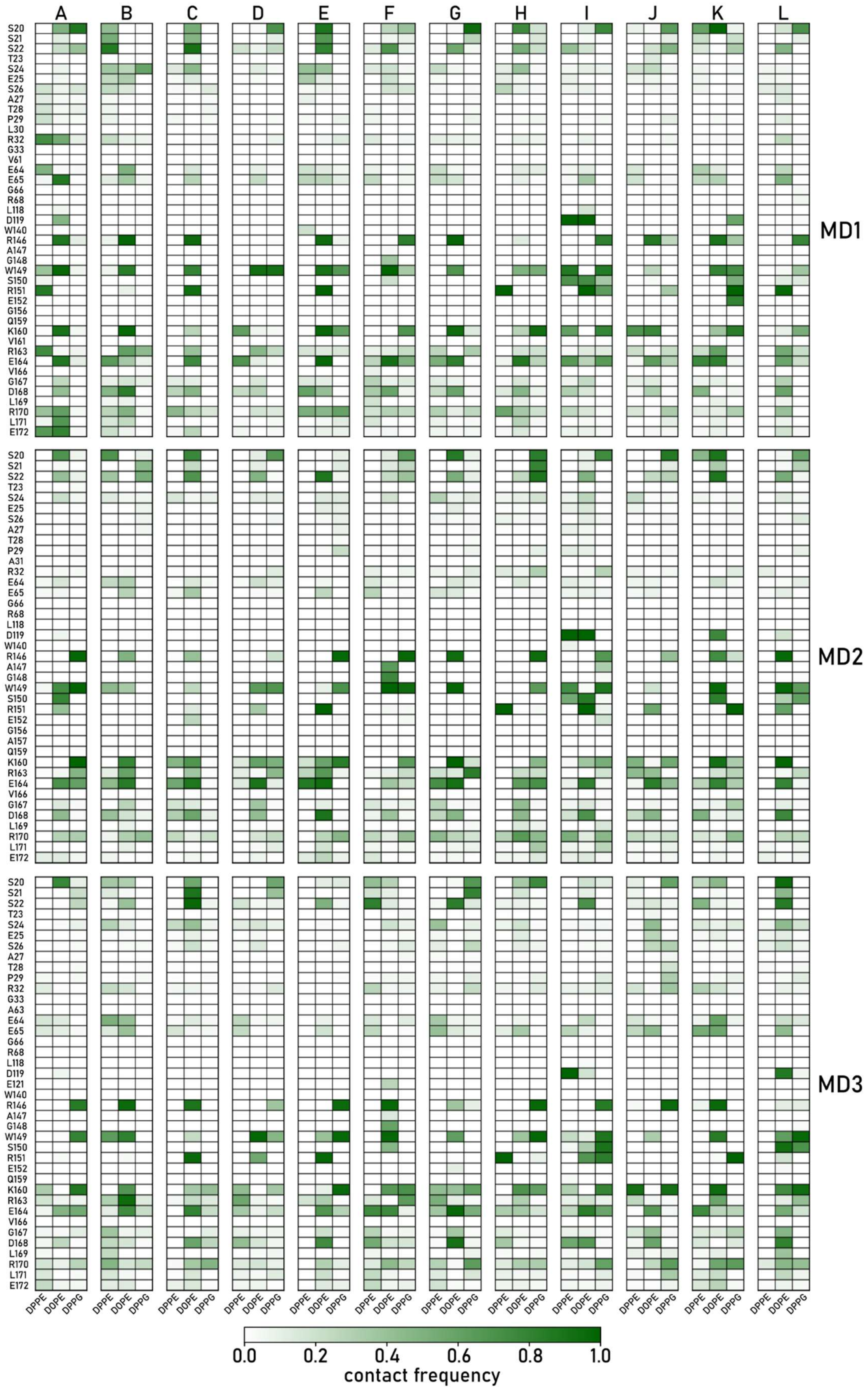


**Supplemental Figure 17. Contact frequency plot of the PelC residues with the lipids of the periplasmic leaflet for each PelC subunit observed in individual simulations.**

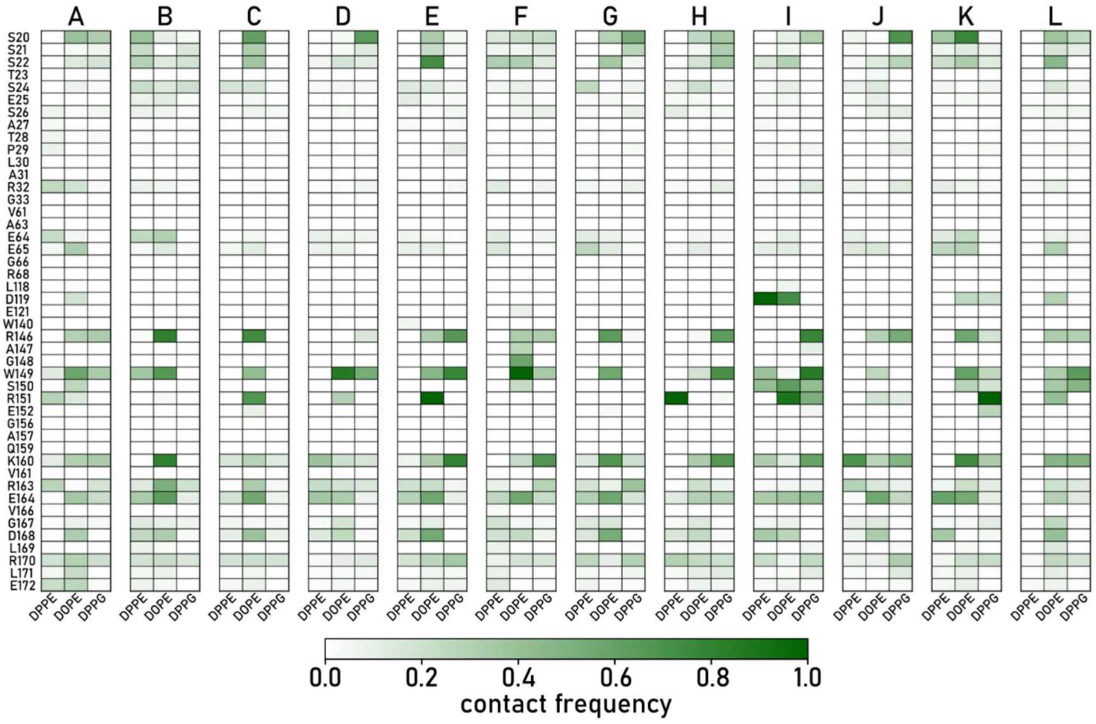


**Supplemental Figure 18. Mean contact frequency plot of the PelC residues with the lipids of the periplasmic leaflet for each PelC subunit, over all three simulations combined.**

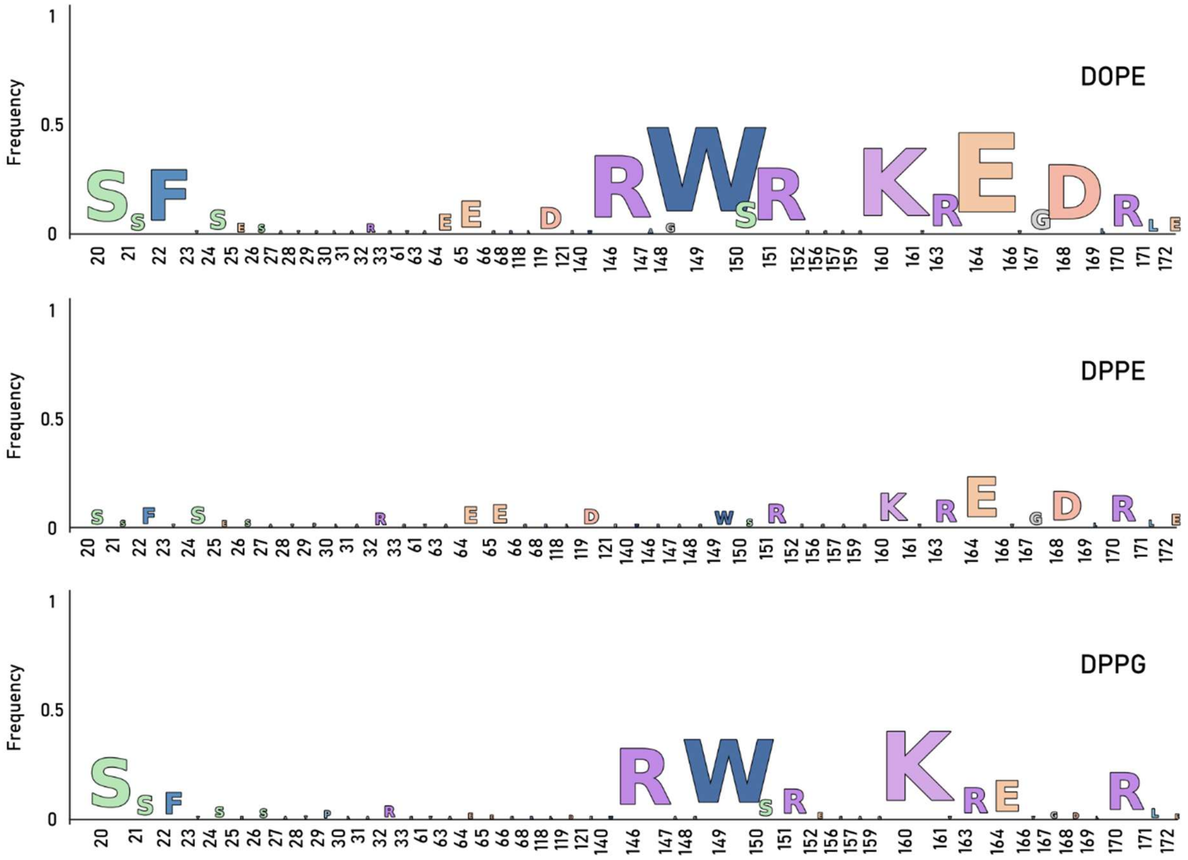


**Supplemental Figure 19. Mean contact frequency of the PelC residues with the inner leaflet lipids presented as a sequence logo over all three simulations combined.**
